# Structural basis for repurposing a flexible phage tail into an Intraspecific bacterial competition weapon

**DOI:** 10.64898/2026.02.24.707460

**Authors:** Fulin Yang, Zelin Zhang, Chengkai Yang, Jun Hou, Hanzhong Feng, Dongsheng Lei, Yong-Xing He

**Affiliations:** School of Life Sciences, Lanzhou University, Lanzhou, 730000, China; School of Physical Science and Technology, Lanzhou University, Lanzhou 730000, People’s Republic of China; State Key Laboratory of Applied Organic Chemistry, Lanzhou University, Lanzhou 730000, China

## Abstract

The rise of multidrug-resistant bacteria poses a major threat to human health. Tailocins, which are phage tail-like bacteriocins, offer a way to kill specific bacteria without the risks of horizontal gene transfer found in phage therapy. While the mechanism of contractile R-type tailocins is well-established, the bactericidal action of non-contractile F-type tailocins remains unresolved. Here, we report the high-resolution cryo-EM structure of the F-pyocin from *Pseudomonas aeruginosa*. The structure reveals a complex nanomachine made of a terminator cap (AlpD/PA0910), a helical tube (PA0633), and a tail tip that mediates a symmetry transition from sixfold to threefold. Notably, the tail tip connects to an elongated central fiber (PA0641) decorated with trimeric side fibers (PA0643/PA0646) involved in host recognition. Through genetic deletion studies and proteomic analysis, we delineate the F-pyocin assembly pathway, establishing that the tape-measure protein (PA0636) serves as a central scaffold coordinating the assembly of the tube, cap, and tail-tip modules. Structural analysis reveals a localized register shift in the central fiber’s coiled-coil shaft, suggesting a metastable state that may facilitate conformational changes upon host recognition. Our findings detail the atomic structure of an F-type tailocin, uncovering its coordinated assembly pathway and specialized tail-fiber architecture, which establishes a structural framework for engineering next-generation, precision antimicrobials.

## Introduction

The rapid emergence of multidrug-resistant (MDR) bacteria represents one of the most urgent threats to global health^1,2^, undermining the efficacy of existing antibiotics and leaving clinicians with limited therapeutic options. Among alternative strategies, phage therapy has shown promise in eradicating otherwise recalcitrant infections^3^. However, concerns remain that therapeutic phages may unintentionally transfer virulence or antibiotic-resistance genes to host bacteria, thereby limiting their clinical application^4^. Tailocins, phage tail–like bacteriocins naturally produced by bacteria to eliminate competitors, provide a strategy to avoid these risks^5^. Unlike phages, tailocins lack a DNA-containing capsid and associated genome packaging machinery, eliminating the potential for horizontal gene transfer^6^. Building on these safety features and their adaptable structure, recent engineering efforts have expanded tailocin targeting to species such as Escherichia coli and Yersinia pestis^7,8^, further highlighting their therapeutic potential.

Tailocins are broadly classified into two groups, R-type and F-type^9^. R-type tailocins resemble myophage tails, employing sheath contraction to puncture target membranes and dissipate the proton motive force^9,10^. In contrast, F-type tailocins consist of flexible, non-contractile tails structurally similar to siphophages, which lack a surrounding sheath^9^. Such tailocin systems are widespread among both Gram-negative and Gram-positive bacteria^11^. For instance, R-type tailocins include the R-pyocin of the opportunistic pathogen *P. aeruginosa*^9,10^ and the diffocin of Clostridioides difficile^11^. Beyond their structural diversity, tailocins are generally considered to play a critical role in kin selection. This ecological function is exemplified by the tailocin from the plant pathogen Pseudomonas viridiflava, which selectively kills genetically similar but non-identical strains while exhibiting little or no activity against unrelated plant-associated bacteria^12^.

Recent advances in cryo-electron microscopy (cryo-EM) have transformed structural studies of phage tail–like systems. For contractile R-type tailocins, atomic models reveal a central core tube encased by a contractile sheath, with a baseplate at one end containing receptor-binding proteins (RBPs)^13^. Binding of an RBP to its receptor triggers baseplate rearrangement and subsequent sheath contraction. This contraction converts stored chemical energy into mechanical force, driving the inner tube through the cell membrane and causing bacterial death^13^. In contrast, the structural details of non-contractile tails have only recently been resolved, primarily due to technical challenges associated with their intrinsic flexibility^11^. To date, among F-type tailocins, only the monocin from Listeria monocytogenes, which resembles the tail structures of TP901-1-like siphophages, has been structurally characterized, revealing a baseplate equipped with three side fibers for receptor binding^11^. Despite these advances, the molecular mechanisms governing the assembly and bactericidal activity of F-type tailocins remain poorly understood^11^.

*P. aeruginosa* is a Gram-negative opportunistic pathogen that poses a significant global health threat due to its intrinsic antibiotic resistance and propensity to cause fatal infections in immunocompromised individuals, such as those with cystic fibrosis, burns, or cancer. It encodes both R-type and F-type tailocins, termed R- and F-pyocins, with their expression induced under conditions that activate the SOS response, such as exposure to DNA-damaging agents like mitomycin C (MMC)^14^. These pyocins are encoded within a single gene cluster located between the trpE and trpG genes^15^. Notably, the F-pyocin exhibits structural features resembling those of λ-like phage tails^15^. Genomic mining of 2,119 *P. aeruginosa* isolates further revealed 11 different groups of F-pyocins, classified based on the specificity modules located at the 3′ end of the clusters^16^. Subsequent functional characterization of six groups demonstrated strain-specific killing patterns that correlate with the LPS O-antigen type of target cells, providing strong evidence that LPS serves as the primary receptor for F-pyocin^16^.

Here, we present the high-resolution cryo-EM structures of F-pyocin from *P. aeruginosa*, providing an atomic-level view of its complex structural organization. By integrating structural analysis with systematic genetic screening and proteomic profiling, we elucidate the fundamental assembly mechanism of this F-type tailocin. Our findings define the principles governing the biogenesis of F-type pyocins and establish a structural framework for understanding how these non-contractile machineries are specialized for bacterial killing.

## Results

### Overall architecture of the F-pyocin

To isolate the F-pyocin machinery for structural analysis, we engineered a *P. aeruginosa* PAO1 derivative strain lacking the genes for the contractile R-pyocin (Δ*PA0622*Δ*PA0623*, hereafter referred to as the parental strain), effectively eliminating contamination from R-pyocins. Following induction with mitomycin C, F-pyocin particles were recovered from the supernatant via PEG 8000 precipitation and further purified by size-exclusion chromatography on a Sepharose 6FF column to yield a homogeneous population.

We determined the atomic structure of the intact F-pyocin assembly by reconstructing four overlapping modules: the cap, helical tube, tail tip, and tail fiber complex (Figure 1B). These modules achieved resolutions of 2.66 Å, 2.60 Å, 2.52 Å, and 3.00 Å, respectively, enabling precise side-chain assignment and atomic modeling of the majority of the structure (SI Appendix, Table S1, SI Appendix, Figures S1 and S2). The resulting composite structure reveals a 145 nm long nanomachine comprising four modular portions, including a cap (Figure 1D), a flexible helical tube (Figure 1D), a symmetry-adapting tail tip (Figure 1G), and a rigid tail fiber complex (Figure 1F). The tube consists of 21 stacked hexameric rings of the tail-tube protein PA0633, forming a continuous channel with an outer diameter of ∼11.5 nm and a lumen of ∼4.0 nm (Figures 1C and 1D). Notably, the density for the proximal cap could not be assigned to any gene within the F-pyocin cluster (*PA0633*–*PA0648*). Through proteomic analysis of the purified particles, we identified the cap as PA0910 (also known as AlpD). AlpD is encoded by the distinct alp operon, which is a system activated by DNA damage to mediate programmed bacterial death (Figure 1A)^17^. The AlpD hexamer seals the proximal end of the PA0633 tube (Figures 1C and 1D), suggesting that *P. aeruginosa* has functionally coupled the assembly of this killing weapon with the host’s suicide-lysis response.

**Figure 1.**
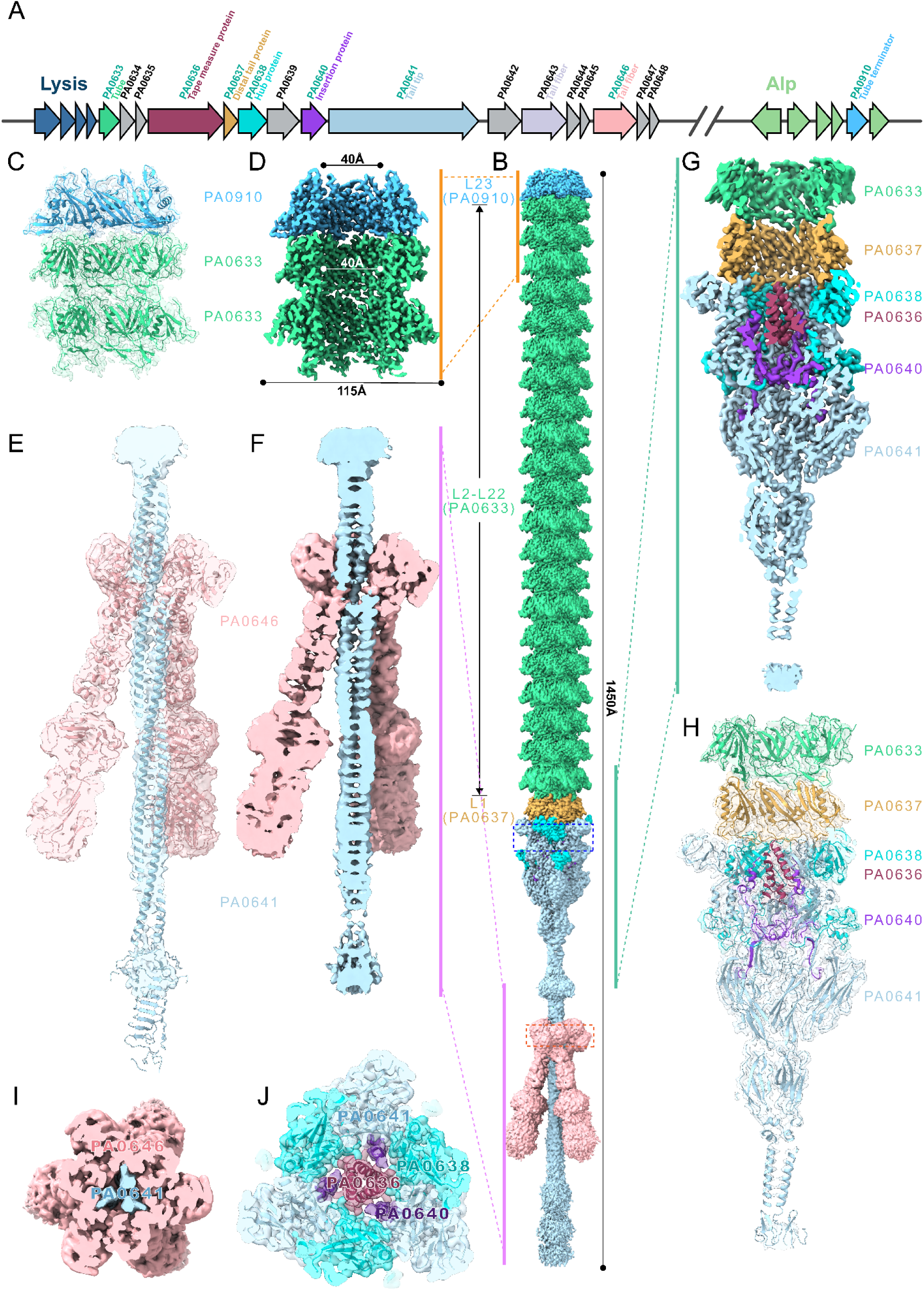
Overall Structure of F-pyocin Particles. **A**, Genomic organization of F-pyocin genes. **B**, Cryo-EM structure of the intact F-pyocin. F-pyocin subunits are color coded and labeled. L1-L23 denotes layers of cap-tube proteins. **C,** Atomic model of the cap (PA0910) and tube (PA0633) fitted into the density. **D,** Cross-sectional view of the cap-tube density, showing its dimensions (115 x 40 Å). **E,** Atomic model of the tail-tip (PA0641) and side-fiber (PA0646) complex. **F,** Cross-section of the tail-tip and side-fiber density, with the PA0633 L2-L22 region emphasized. **G,** Cross-sectional view detailing the sequential, layered assembly of tail proteins. **H,** Corresponding atomic model of the tail-tip complex. **I,** Orthogonal cross-section (90°) revealing the trimeric organization of PA0646. **J,** Top view of the PA0636-PA0641 subcomplex, depicting the central interface for complex assembly.

At the distal end, the transition from the six-fold symmetry of the tube to the three-fold symmetry of the killing apparatus is mediated by the distal tail protein PA0637. A hexamer of PA0637 attaches immediately beneath the terminal tube ring, docking onto a complex formed by trimers of the hub protein PA0638 and the central fiber protein PA0641 (Figures 1G and 1H). This assembly seals the distal lumen, which encloses the C-terminal coiled-coil region of the tape-measure protein (TMP, PA0636, residues 579–611) ^18^ (Figure 1J).

Extending distally from the tip complex is a long, fine central fiber formed by the C-terminus of PA0641 (Figures 1E and 1F). This central fiber serves as the scaffold for host recognition and is decorated by three side fibers attached via their N-terminal domains to a triple-helix bundle on the central fiber shaft (Figure 1I). Although the N-terminal domains (residues 2–76) of the two fiber protein isoforms, PA0643 and PA0646, share ∼90% sequence identity, mass spectrometry of our purified samples revealed a significantly higher abundance of PA0646. Consequently, we modeled the resolved side-fiber density as PA0646. Notably, the putative chaperones encoded downstream of the fiber genes (*PA0644*/*PA0645* and *PA0647*/*PA0648*) remained undetectable in the proteomic profile of the purified particles, indicating that these side fibers represent the mature form of the apparatus.

### AlpD capping the F-pyocin tube is dispensable for bactericidal activity

The F-pyocin tail tube is composed of hexameric rings of PA0633 arranged in a right-handed helix with a rise of ∼40 Å and a twist of 22.1° (Figures 2A and 2B). Each PA0633 subunit adopts a typical siphophage tail protein fold^18^, featuring a central β-sandwich domain and an extended β-hairpin loop that interlocks with adjacent subunits to stabilize the assembly (Figure 2D). The central channel of the tail tube exhibits a strong negative electrostatic potential (Figures 2E and 2F). The tube is capped by a hexamer of AlpD, which shares the fold of the tube protein PA0633 but lacks the outer β-sheet interface required for further polymerization (Figures 2H, 2I and 2G). This structural distinction prevents the addition of subsequent subunits, effectively terminating tube elongation. The AlpD hexamer also displays a distinct Ω-shaped loop that protrudes into the channel lumen, restricting the inner diameter to 4.0 nm (Figure 2I). As AlpD is a component of the *P. aeruginosa* alp operon^17^, its identification as the F-pyocin cap reinforces the model of functional coupling between tailocin assembly and the Alp-dependent cell lysis required for its release^17^.

**Figure 2.**
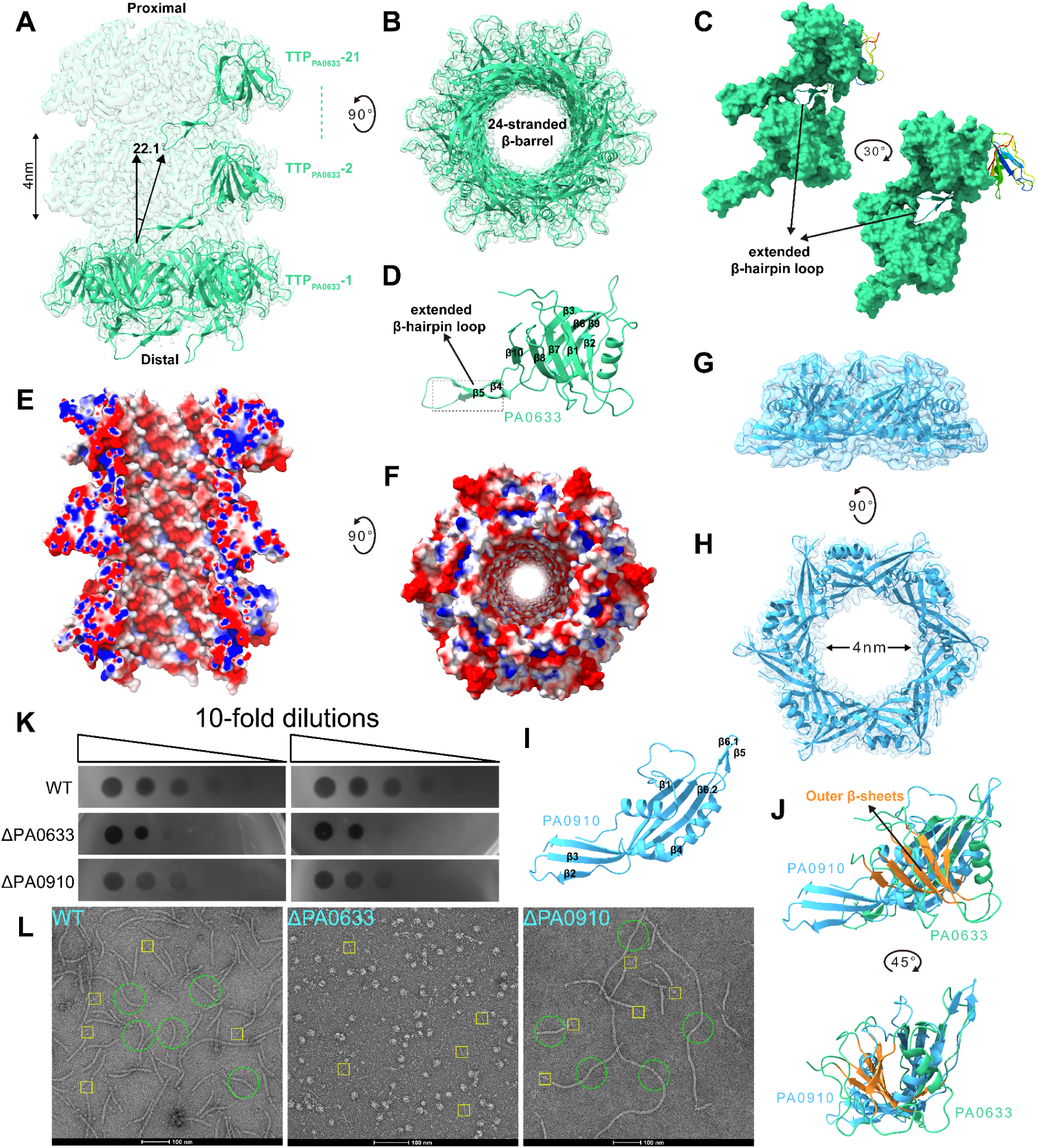
Architecture of the F-pyocin tube and cap complexes. **A,** Overall tube architecture, with 21 stacked layers (4 nm layer height, 22.1° rotation between layers). **B,** Top-down view showing the 24-stranded β-barrel. **C,** Close-up of the assembly interface, highlighting the extended β-hairpin loop that mediates inter-layer stacking. **D,** PA0633 tube subunit with its extended β-hairpin loop. **E,** Electrostatic surface potential of the tube. **F,** Same view rotated 90°, showing inner and outer surface charge distribution. **G,** Overall structure of the cap complex. **H,** Cap rotated 90°, revealing a 4 nm central channel. **I,** Ribbon diagram of a single PA0910 cap subunit. **J,** Structural overlay of cap (PA0910) and tube (PA0633) monomers—note the additional outer β-sheet in the tube subunit. **K,** Functional assays. Spot titrations comparing bactericidal activity of WT, Δ*PA0633*, and Δ*PA0910* mutants. **L,** Corresponding negative-stain micrographs showing particle integrity from these mutants.

To evaluate the roles of PA0633 and AlpD, we generated deletion strains (Δ*PA0633* and Δ*AlpD*) in the background of the parental strain and analyzed their lysates by negative-stain electron microscopy (nsEM). The Δ*PA0633* strain failed to assemble F-pyocins (Figure 2L), resulting in a marked loss of bactericidal activity (Figure 2K). In contrast, the Δ*AlpD* strain produced F-pyocin particles with intact tail tips but exceptionally long, unregulated tail tubes (Figure 2L). This phenotype confirms that AlpD functions as the terminator of tube polymerization. Interestingly, the Δ*AlpD* mutant exhibited bactericidal activity almost comparable to that of the wild-type (Figure 2K). This observation indicates that while AlpD is essential for precise length control, it is not strictly required for the bactericidal function of the F-pyocin^17^.

### Structural basis of tail tip assembly and the proteolytic checkpoint

The F-pyocin tail tip constitutes an inverted cone-like apparatus connecting the hexameric tube to the trimeric tail fibers (Figure 3A). At the tube-tip junction, the distal tail protein PA0637 forms a hexameric ring that latches onto the protruding β-hairpin arms of the terminal tube hexamer, thereby terminating tube polymerization on the proximal side. This hexameric ring sits directly atop the trimeric hub core, which exhibits pseudo-hexameric symmetry and is co-assembled by the tail hub protein PA0638 and the N-terminal domain of PA0641 (residues 373–594) (Figures 3C and 3D). To accommodate the symmetry transition from the hexameric PA0637 adapter to the trimeric hub, each tail-tube-fold-like domain from the heterodimeric PA0638/PA0641 unit interacts with a dimer of PA0637 (Figure 3A). This 6-to-3 symmetry mismatch is resolved at the interface where the long protruding loops of PA0637 anchor onto the upper surface of the PA0638/PA0641 ring. Notably, each PA0638 subunit coordinates a 4Fe-4S cluster at the interface with PA0641 via four conserved cysteine residues, resembling the redox-active center in the hub protein gpL of bacteriophage λ^19^ and likely acting as a “molecular pin” to maintain the structural integrity of the killing apparatus (Figures 3E and 3H).

**Figure 3.**
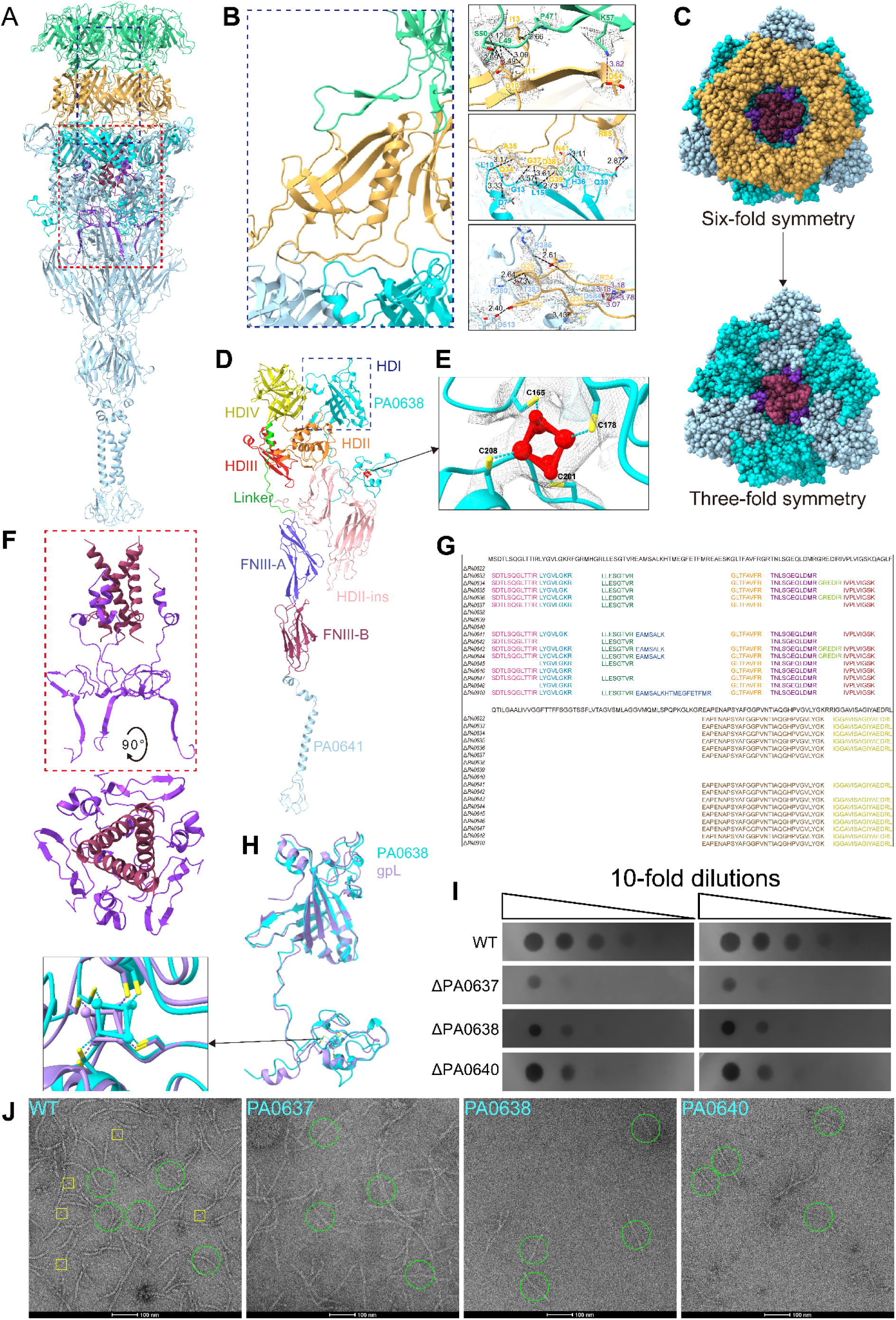
Structural architecture and symmetry transition of the F-pyocin tail-tip complex. **A,** Overall architecture of the tail-tip complex. **B,** Key interfaces mediating PA0633–PA0637, PA0637–PA0638, and PA0637–PA0641 assembly, showing how an extended E-loop anchors hexameric PA0637 to trimeric PA0638/PA0641. **C,** Schematic of the 6-to-3 symmetry mismatch critical for tail-tip integrity. **D,** Domain organization of PA0641, including HD (Ⅱ-IV), linker, and FNIII modules. **E,** The [4Fe-4S] cluster in PA0638 with coordinating cysteines (C165, C178, C201, C208). **F,** Orthogonal views of the PA0636–PA0640 subcomplex. **G,** Peptide coverage of PA0640 across various F-pyocin deletion mutants. **H,** Structural alignment showing PA0638 as a conserved “molecular pin” akin to λ phage gpL. **I,** Spot titration assays comparing bactericidal activity of WT, Δ*PA0637*, Δ*PA0638* and Δ*PA0640* mutants. **J,** Negative-stain micrographs of particles assembled from these mutants, revealing effects on structural integrity.

The largest component, PA0641, contains two distinct fibronectin type III (FNIII) domains that seal the distal end of the tail tip (Figure 3D). Immediately downstream of these FNIII domains, the cryo-EM map resolved a trimeric α-helix bundle, which transitions into a distal region predicted by AlphaFold3 to adopt a β-prism fold^20^ (Figures 4A and 4F). The internal lumen of the tail tip is occluded by a trimer of PA0640, a structural homolog of λ phage gpI^21^, which we termed the tip plug protein (Figure 3F). Notably, only the C-terminal portion (residues 127–200) of the tip plug is visible in the electron density map, suggesting its N-terminal region is either highly flexible or proteolytically cleaved (Figure 3F). The N-terminal α-helices of this plug protein encircle the C-terminal 33 residues (579–611) of the TMP (PA0636), which adopts a coiled-coil conformation (Figure 3F). Concurrently, the plug protein anchors itself by interacting with multiple regions of PA0638 and PA0641 via its C-terminal loops, effectively bridging the TMP to the tail tip. This arrangement likely allows the plug to both recruit the TMP for tail length determination and lock the tail tip in a closed state, thereby preventing premature TMP release before host recognition.

**Figure 4.**
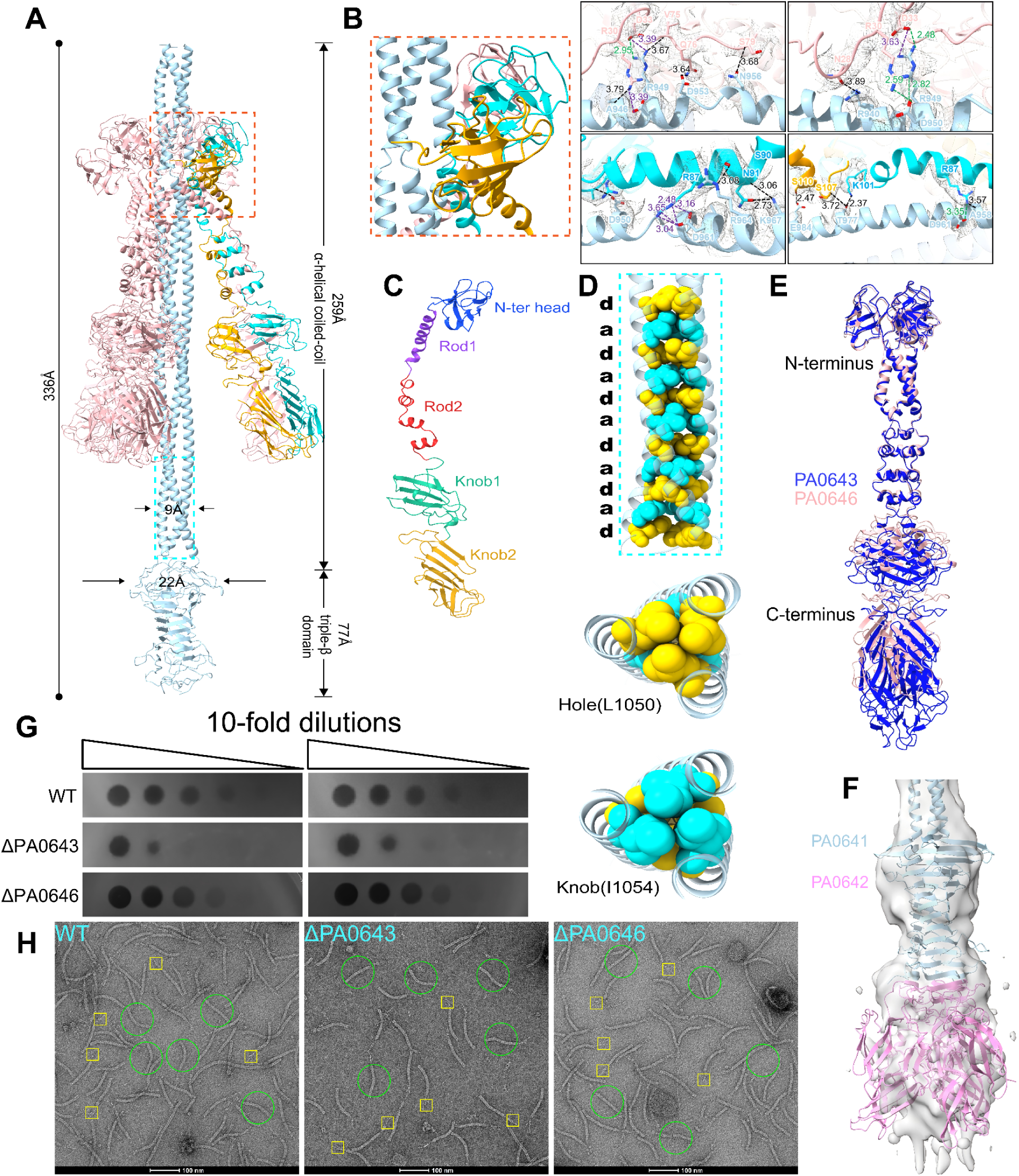
Organization and host-recognition logic of the F-pyocin tail-fiber complex. **A,** Overall structure of the tail-fiber region, with the PA0646 fiber highlighted. **B,** Close-up view of the side-fiber anchoring interface, detailing interactions between PA0646 and PA0641. **C,** Modular domain organization of the tail-fiber components (Rod1, Rod2, Knob1, Knob2). **D,** Trimeric assembly interface, with key oligomerization residues L1050 (“hole”) and I1054 (“knob”) highlighted. **E,** Structural comparison (superposition) of the VIA-type specificity modules PA0643 and PA0646. **F,** Superimposition of the AlphaFold3-predicted model (distal β-prism domain and PA0642) with the cryo-EM density, showing high consistency. **G,** Host-range assays. Spot titrations showing bacteriolytic activity of WT, Δ*PA0643* and Δ*PA0646* mutants against P. aeruginosa clinical isolates. **H,** Negative-stain EM micrographs showing morphological changes of P. aeruginosa cells treated with WT and mutant F-pyocin variants.

To dissect the functional roles of the tail tip components, we generated single deletion mutants of the tip components (Δ*PA0637*, Δ*PA0638*, and Δ*PA0640*) and the TMP (Δ*PA0636*). Examination of lysates from these mutants by nsEM revealed tail tube structures completely lacking tail tips and fibers (Figure 3J), leading to a marked reduction in bactericidal activity (Figure 3I). Intriguingly, mass spectrometry analysis revealed a distinct stability pattern where the internal plug protein PA0640 was undetectable in the Δ*PA0638* mutant, indicating that PA0640 requires the hub protein for recruitment or stability. In contrast, PA0640 was detected in the Δ*PA0637* and Δ*PA0636* backgrounds, suggesting that the hub core (comprising PA0638 and PA0640) can assemble but fails to dock onto the tail tube in the absence of the adapter PA0637 or TMP PA0636 (Figure 3G). Morphological analysis by nsEM further highlighted the regulatory role of the adapter, as the Δ*PA0637* mutant exhibited a unique phenotype of exceptionally long, unregulated tail tubes reminiscent of the Δ*AlpD* cap mutant, whereas other mutants displayed heterogeneous tube lengths without extensive long tube formation. These data collectively indicate that PA0637 is essential for restricting tube polymerization at the distal end, while the concerted action of the complete tail tip and TMP is required for precise length determination.

Careful inspection of the mass spectrometry data for mature wild-type F-pyocin revealed that the N-terminal region of the tail plug protein PA0640 was undetectable, consistent with the missing density in our cryo-EM structure (Figure 3G). To distinguish between low ionization efficiency and genuine proteolytic processing, we analyzed the Δ*PA0633*, Δ*PA0637*, and Δ*PA0636* mutants, in which the tail tip complex assembles but fails to attach to the tail tube. In these F-pyocin assembly-stalled backgrounds, the N-terminal region of PA0640 was detected, confirming its presence in the precursor tip.

Furthermore, when full-length PA0640 was recombinantly expressed alone in *E. coli*, the N-terminal region was spontaneously removed by unknown host proteases, suggesting its intrinsic instability (SI Appendix, Figure S3). Collectively, these results indicate that while the N-terminal domain is protected from degradation within the assembled premature tail tip, it must undergo specific proteolytic removal to facilitate the attachment of the tail tip to the tail tube and generate the mature F-pyocin particles^22,23^.

### Organization of the tail fiber complex

The C-terminal region of PA0641 extends from the tail tip to form a long, trimeric coiled-coil shaft that serves as the binding platform for the trimeric side tail fibers (PA0643 and PA0646) (Figure 4A). While the cryo-EM map resolves only the coiled-coil region of PA0641 (residues 907–1084), AlphaFold3 predictions^20^, supported by our proteomics data, indicate that the distal end adopts a β-prism fold to which a trimeric PA0642 complex attaches in the mature F-pyocin (Figure 4F). Structural analysis using SOCKET2 confirmed that this shaft domain (residues 913–1085) predominantly adopts a typical trimeric coiled-coil architecture characterized by a continuous heptad repeat register (abcdefg)_n_ (Figure 4D)^24^. The C-terminal segment (residues 1050–1085) exhibits highly stable “knobs-into-holes” packing^24,25^, where the hydrophobic core at the ‘a’ and ‘d’ positions is supplemented by polar residues that likely stabilize the trimeric interface through localized hydrogen-bond networks (Figure 4D). However, this long coiled-coil is punctuated by a significant structural “stutter” near the C-terminus (residues 1070–1082). Register analysis reveals a transition from strict heptad periodicity to a localized register shift, marked by a ‘defg’ pattern jump^24^. This discontinuity induces a localized under-winding of the superhelix^25^, resulting in a slightly expanded core with reduced packing density. Strategically positioned upstream of the predicted β-prism domain, this “mechanical weak point” likely serves as a conformational hinge or nucleation site. Under mechanical shear, this stutter may facilitate the transition from the coiled-coil state to an extended β-structure by lowering the energetic barrier for helical unwinding.

Three side tail fibers dock around this kinked region of the coiled-coil shaft with pseudo-C3 symmetry (Figure 1I), with each fiber trimer utilizing the N-terminal domains of two of its protomers to attach to the central shaft(Figure 4B). Sequence analysis reveals that the N-terminal attachment regions of the two side fiber isoforms, PA0643 and PA0646, share 80% sequence identity. In contrast, their distal ends share only 30% identity and structurally resemble the C-terminal fragments of R-pyocin fibers^26^, implying that these two sets of side fibers recognize different host receptors.

To investigate the specific functions of the central and side fibers, we constructed deletion mutants lacking the PA0641 C-terminus (residues 1051–1255), PA0642,PA0643, or PA0646. In the Δ*PA0641* and Δ*PA0642* mutants, although tail fibers were still observed, the purified F-pyocins displayed heterogeneous tube lengths and a marked reduction in bactericidal activity (Figure 4J). Conversely, the particles produced by the Δ*PA0643* and Δ*PA0646* mutants were morphologically similar to wild-type F-pyocins(Figure 4J). This suggests that the distinct fiber isoforms encoded by *PA0643* and *PA0646* likely assemble into homotrimers in a mutually exclusive manner^26^, equipping subpopulations of F-pyocins with varying binding specificities to broaden the target spectrum^27^. Deletion of PA0643 led to a significant reduction in bactericidal activity against our tested strain, whereas deletion of PA0646 had little effect, strongly suggesting that these two sets of tail fibers are involved in recognizing different host receptors^27^ (Figure 4I).

### Molecular chaperones during F-pyocin assembly

Phage tail assembly is a highly coordinated process that strictly relies on various transient chaperone proteins^28,29^. Consistent with this observation, several genes within the F-pyocin operon, namely PA0634, PA0635, PA0639, PA0644, PA0645, PA0647, and PA0648, are absent from our cryo-EM structure and undetected in the proteomic data of mature F-pyocin particles (SI Appendix, Figure S4), suggesting they function as assembly factors. Based on genomic context (Figure 1A), PA0634 and PA0635 are predicted to assist in tail polymerization, while PA0639 shares homology with the λ phage tail tip assembly protein gpK. Additionally, PA0644/PA0645 and PA0647/PA0648 are predicted to encode specific tail fiber assembly chaperones (TFACs) dedicated to PA0643 and PA0646, respectively^30^.

To systematically investigate these putative assembly components, we constructed single deletion mutants and evaluated them using nsEM and bactericidal assays. The Δ*PA0634* mutant exhibited a phenotype identical to the Δ*PA0636* (TMP) mutant, producing abnormal, heterogeneous tail tubes devoid of tail tips and fibers (Figure 5A), alongside a complete loss of bactericidal activity (Figure 5B). This suggests that PA0634 is an indispensable chaperone for the stabilization or proper functional loading of the TMP during initial assembly^18^. Strikingly, the Δ*PA0635* mutant resembled the phenotype of the Δ*PA0637* adapter mutant, forming exceptionally long, unregulated tubes lacking the tip apparatus (Figure 5A), which strongly indicates that PA0635 is specifically required for the correct folding or recruitment of the PA0637 hexamer, as its absence prevents the adapter from capping the growing tube and results in uncontrolled distal polymerization^31^. Furthermore, the Δ*PA0639* mutant displayed a phenotype closely resembling the Δ*PA0638* and Δ*PA0640* hub complex mutants, generating non-functional, tip-less tubes (Figure 5A). This implies that PA0639 functions as a critical maturation factor essential for the assembly or structural integrity of the tail tip complex.

**Figure 5.**
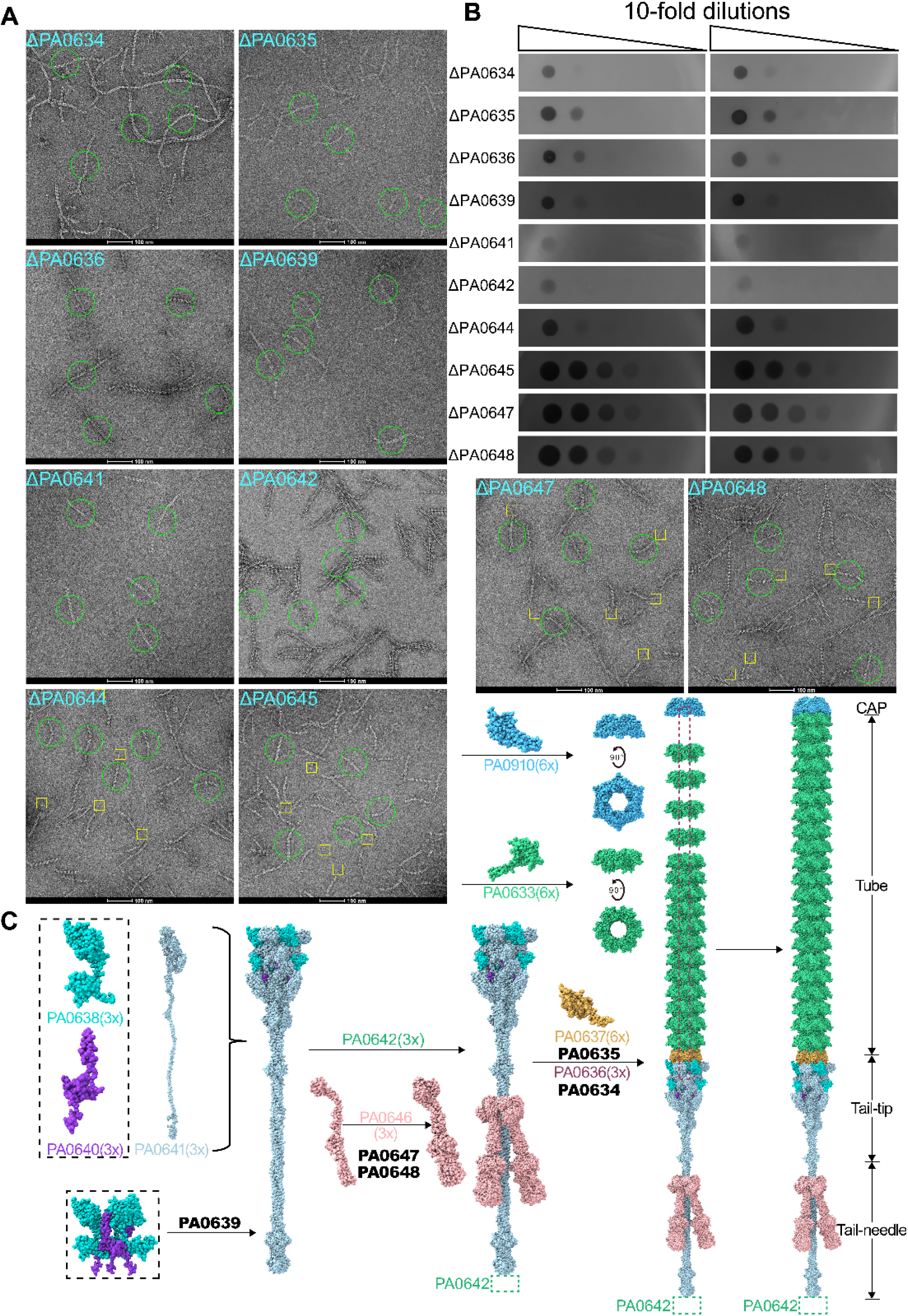
Assembly mode analysis of F-pyocin. **A, B,** Host-range and bactericidal spectrum assays. A, Spot titration assays showing the bacteriolytic activity of Δ*PA0634*, Δ*PA0635*, Δ*PA0636*, Δ*PA0639*, Δ*PA0641*, Δ*PA0642*, Δ*PA0644*, Δ*PA0645*, Δ*PA0647* and Δ*PA0648* single deletion mutants against a panel of *P. aeruginosa* clinical isolates. B, Negative-stain electron micrographs illustrating the morphological changes of *P. aeruginosa* cells upon treatment with Δ*PA0634*, Δ*PA0635*, Δ*PA0636*, Δ*PA0639*, Δ*PA0641*, Δ*PA0642*, Δ*PA0644*, Δ*PA0645*, Δ*PA0647* and Δ*PA0648* single deletion mutants. **C,** F-pyocin assembly mode diagram.

Unexpectedly, evaluation of the putative TFACs revealed a functional divergence within the PA0644/PA0645 pair. The Δ*PA0644* mutant exhibited a complete loss of bactericidal activity (Figure 5B), resembling the severe defect of the Δ*PA0633* major tail tube mutant. In contrast, the Δ*PA0645* mutant retained wild-type levels of activity (Figure 5B), suggesting that PA0645 is functionally dispensable or its role is compensated by other factors^32,33^.

These findings suggest an assembly model for F-pyocin where the process begins at the distal tail tip with the formation of the trimeric hub core, comprising PA0638, PA0640 and PA0641, under the guidance of the maturation factor PA0639^31^ (Figure 5C). Since PA0641 is an integral component of the hub core, the central fiber is established during this initial stage, which also includes the attachment of PA0642 and the side fibers, PA0643 and PA0646, facilitated by chaperones such as PA0644/PA0645 and PA0647/PA0648 (Figure 5C). Subsequently, the hexameric adapter PA0637, whose folding depends on PA0635, docks onto the trimeric hub to resolve the symmetry mismatch, while the TMP, stabilized by PA0634, is anchored into the hub-plug complex (Figure 5C)^28^. This complete complex then nucleates the major tail tube protein PA0633, which polymerizes upward along the TMP scaffold until extension is terminated at the proximal end by the cap protein AlpD to ensure a precisely determined length (Figure 5C)^28^. This coordinated assembly process results in structurally intact F-pyocins, with the TMP sequestered by the internal plug protein PA0640 until host recognition triggers penetration^34^.

## Discussion

In this study, we elucidate the complete, near-atomic structure and assembly pathway of the F-type pyocin from *P. aeruginosa*. Our findings provide the first detailed molecular insight into this class of non-contractile tailocins involved in interspecies competition. A defining feature identified in our work is the fine architecture of the tail fiber, which consists of an extended central fiber anchored by three side fibers. The majority of the central fiber is a coiled-coil shaft terminated by a globular domain that binds to PA0642, a putative adhesin responsible for recognizing specific receptors on target strains. Structural analysis of the coiled-coil shaft revealed that while most regions exhibit stable “knobs-into-holes” packing^35^, the distal region is punctuated by a discontinuity in its hydrophobic core. This localized instability disrupts the ideal coiled-coil geometry, thereby lowering the energetic barrier for unwinding. Notably, during the preparation of this manuscript, a complementary study by He et al., recently appearing as a preprint, also reported the structure of the F-pyocin and captured the post-ejection state of the tail tip adopting a stable β-prism fold. Taken together, these results suggest that the metastable coiled-coil shaft serves as a predesigned energy reservoir essential for the ejection of internal proteins and the binding of PA0642 to its cognate receptor may trigger a concerted conformational change^36,37^, leading to the unwinding of the coiled coil shaft and the subsequent ejection of the tape measure protein.

Our data also reveal that F-pyocin assembly is a highly orchestrated process, regulated by a proteolytic checkpoint and a suite of dedicated chaperones. We found that the N-terminal domain of the tip plug protein, PA0640, is proteolytically removed prior to the attachment of the tail tip to the tail tube. This suggests that the unprocessed N-terminal domain may act as a steric shield, preventing premature docking of the tail tip to the tube and the tape measure protein until the tip is fully matured^38^. Our genetic and structural analyses further reveal that PA0649 is essential for tail tip assembly, while PA0645 functions as a specific chaperone for the distal tip protein PA0647, likely to prevent its premature aggregation^18^. However, the exact molecular mechanisms governing the cleavage of PA0640 and the precise coordination of these chaperones in F-pyocin assembly remain to be fully elucidated.

The integration of AlpD as the hexameric cap protein underscores a functional cross-talk between F-pyocin assembly and the alp-mediated programmed cell death (PCD) pathway^17^. Although the alp operon encodes a potent holin (AlpB) that forms inner-membrane pores and triggers autolysis in *P. aeruginosa*, it lacks its own endolysin and spanin genes^39^, which are required for complete cell lysis. This genomic organization strongly suggests that alp-mediated PCD is functionally dependent on the F-pyocin locus, which provides the necessary endolysin and spannins to finalize the lysis process. While the pyocin operon contains its own lysis cassette^15^, the alp system likely provides a redundant yet high-potency holin via AlpB, ensuring the robustness of the lysis-dependent release of F-pyocins. Furthermore, by utilizing AlpD as a cap to precisely control the termination of tube polymerization, *P. aeruginosa* can optimize the length of F-pyocin particles, thereby maximizing its metabolic investment in “weapon” production and ensuring the efficient dissemination of mature particles across the bacterial population.

In summary, we have determined the atomic structure and assembly mechanism of the non-contractile F-type pyocin. Distinct from their contractile counterparts, F-type pyocins achieve specific target recognition through an elaborate tail fiber architecture. Furthermore, the identification of AlpD as the hexameric cap protein reveals a functional cross-talk between F-pyocin assembly and the host bacterial programmed cell death machinery, ensuring that F-pyocin production and cell lysis are tightly synchronized. These results establish a comprehensive structural framework for engineering tailocins into a new class of precision antimicrobial agents to combat the growing threat of antibiotic resistance^6^.

## Acknowledgment

We acknowledge the Electron Microscopy Centre of Lanzhou University for assistance with cryo-electron microscopy data collection. We also acknowledge the Mass Spectrometry Facility of the School of Life Sciences at Lanzhou University for support with proteomic data acquisition. This study was supported by the Gansu Research Program (Grant No 22CX8NA001), the National Natural Science Foundation of China (Grant No 31971422), the National Natural Science Foundation of China (Grant No. 32171300)

## Author contributions

Y.-X.H., D.L., Y.W., and F.Y. conceptualized and designed the research; F.Y., C.Y., performed the experiments; Z.Z, performed cryo-EM data collection and structure determination. D.L., J.H., F.Y., H.G., and H.F. contributed to data collection and analysis; F.Y., Z.Z., and Y.-X.H. drafted the manuscript; Y.-X.H. revised the manuscript. All authors have read and approved the final version of the manuscript.

## MATERIAL AND METHODS

### Bacterial strains, plasmids and culture condition

The plasmids, bacterial strains, and primers used in this study are listed in Tables S2, S3, and S4 in the SI Appendix, respectively. Escherichia coli strains BL21 (DE3), DH5α, and WM3604 were cultured at 37 °C in Luria-Bertani (LB) broth. *P. aeruginosa* PAO1 and its derivative strains were grown at 37 °C in LB broth.

### Construction of mutation strains

In-frame deletion mutants were constructed using a two-step homologous recombination method^40^. Approximately 1 kb of the upstream and downstream regions of the target genes was amplified by PCR from *P. aeruginosa* PAO1 genomic DNA using primers listed in Table S4 in the SI Appendix. These fragments were digested with specific restriction enzymes and ligated into the suicide vector pK18mobsacB. The resulting plasmid was transformed into E. coli WM3064 and subsequently introduced into the recipient strain via conjugation. Mutants were selected on media containing 15% sucrose, and successful deletion was confirmed through PCR analysis following established protocols.

### Induction and Purification of pyocins

To generate lysates containing pyocins and phages from the 17 strains, all strains were first streaked out on LB plates and grown at 37 °C for 14 hours. Next, overnight cultures were started by inoculating fresh LB with a single colony and were grown for 14-16 hours at 37 °C. The overnight cultures were used to inoculate 500 mL of fresh LB (1% inoculum was used) and grown at 30 °C until the cells reached a density of OD600 = 0.4. An inducing agent, Mitomycin C, was added to a final concentration of 4 μg/mL and shaking at 30 °C was resumed for 5 hours or till lysis. The induced cultures were cooled to room temperature (RT) and 4 U/mL of DNaseI was added and shaken at 30 °C for 30 min. The bacterial supernatant was supplemented with 0.1% (v/v) Triton X-100, followed by gradual addition of ammonium sulfate to 176 g/L (30% saturation) with continuous stirring at 4 °C overnight. After centrifugation at 4,500 × g for 1 h at 4 °C, the precipitate was resuspended in 10 mL SM buffer (100 mM NaCl, 8 mM MgSO₄, 50 mM Tris-HCl, pH 7.5). Concurrently, the retained supernatant was brought to 60% saturation by adding 188 g/L ammonium sulfate and stirred overnight at 4 °C.The mixture was centrifuged again under identical conditions (4,500 × g, 1 h, 4 °C). The resulting precipitate was resuspended in 10 mL of SM buffer and pooled with the previous 30% saturation fraction. This pooled suspension was diluted to 100 mL with SM buffer, followed by addition of 75 g/L PEG8000 and 30 g/L NaCl under overnight stirring at 4 °C. Post-centrifugation (4,500 × g, 1 h, 4 °C), the pellet was resuspended in 20 mL SM buffer containing 0.1% (w/v) n-Dodecyl β-D-maltoside (DDM) and incubated overnight at 4 °C with agitation. Insoluble material was removed by centrifugation at 13,000 × g for 1 h, and the clarified supernatant was filtered through a sterile 0.22 μm cellulose acetate membrane.The sample was concentrated using a 30 kDa Amicon Ultra centrifugal filter to about 500 μL, and further purified via size-exclusion chromatography on a Superdex 200 10/300 GL column pre-equilibrated with SM buffer (100 mM NaCl, 8 mM MgSO₄, 50 mM Tris-HCl, pH 7.5). Elution profiles were monitored at 220 nm and 280 nm, and fractions containing purified protein were pooled, concentrated again, and stored at −80°C.

### Negative-stain electron microscopy

The negative-stain electron microscopy specimens were prepared as described in a previously published protocol^41^. In brief, a 4 µL aliquot of the sample was applied onto a continuous carbon film grid (CF200-Cu, Electron Microscopy Sciences) that had been glow-discharged. Excess solution on the grid was blotted after ∼1 min incubation. The grid was then washed with three drops of water and stained with three drops of uranyl acetate before drying with an oven lamp. The grid was examined on an Talos F200C (FEI) electron microscope.

### Cryo-EM sample preparation and data collection

A 3.5 μL aliquot of the F-pyocin sample was applied to a glow-discharged Quantifoil 300 mesh Au 1.2/1.3 grid (Quantifoil Micro Tools). The grid was plunge-frozen in liquid ethane using a Vitrobot Mark IV (Thermo Fisher Scientific) with a blot time of 5-10 s, at 4 °C and 100% humidity. Cryo-EM data were acquired on a Thermo Fisher Scientific Titan Krios G3i microscope equipped with a Gatan Bioquantum K3 direct electron detector. Automated data collection was performed using EPU software in super resolution mode, with an exposure time of 3 s over 40 frames, a defocus range of −0.8 to −2.2 μm, and a nominal pixel size of 0.83 Å.

### Cryo-EM data processing

Data processing was carried out in cryoSPARC^42,43^. Movie frames were motion-corrected with patch-based alignment, and CTF parameters were estimated using CTFFIND4^44,45^. An initial set of 100 particles were manually picked out for 2D classification, with representative 2D class averages selected as templates for automated particle picking, yielding 1,896,237 particles. After multiple rounds of 2D classifications, 755,357 high-quality particles were selected and subjected to 3D classification. The best-resolved subset of particles (108,026) for the tail-tip region was re-centered and locally refined to obtain the final density map.

For the cap, tube, and tail fiber regions, Topaz models were trained separately based on their respective 2D class averages, yielding 152,365, 1,261,647, 199,058 particles, respectively. After multiple rounds of 2D classifications, 103,687 (cap), 1,197,486 (tube) and 36,768 (tail-fiber) high-quality particles were selected. Initial low-resolution 3D models were generated and further classified to identify the best subsets: 89,290 particles for the cap, 230,514 for the tube, and 35,179 for the tail-fiber. The cap and tube regions were refined using 3D auto-refinement, while the tail-fiber region was re-centered and locally refined to generate the final density maps. The resolution of all maps was estimated using the gold-standard Fourier shell correlation (FSC) cutoff of 0.143.

### Model building, refinement and validation

Due to structural flexibility, the tail-fiber region exhibited lower resolution toward its tip. Initial models for the cap, tube, and tail-tip regions were generated using ModelAngela^46^, while the tail-fiber model was constructed by integrating ModelAngela and AlphaFold3^20^. All the initial models were submitted to real space refinement in Phenix^47^ and manual adjustments in Coot^48^. Final models were validated using Phenix, and the validation statistics are summarized in Table S1 in the SI Appendix.

### Quantification of pyocin tail length

To quantify the number of layers in the tail tube of F-pyocin, a total of 200 particles from the cryo-electron micrographs were analyzed. The distribution of layer numbers was fitted to a normal distribution, with 21 layers being the most frequent.

### Spot assay

100 μL of overnight culture of each strain was added to 7.5 mL of top agar (0.6% agar), which was immediately plated on a LB plate to create a bacterial lawn. Right after solidification, on each lawn 2 μL of 10-fold serial dilutions of each lysate were spotted. Plates were incubated at 37 °C for 14 hours. Zones of clearing and physical characteristics of each clearing were carefully noted.

### Plasmid Construction

Two full-length tail fiber genes were amplified from the *P. aeruginosa* PAO1 strain. For the construction of the two Tail fiber complex plasmids, PA0643/PA0646 were subcloned into the first cloning site of the pETDuet-1 vector (between the BamHI and HindIII restriction enzyme sites), forming a fusion protein bearing an N-terminal 8×His tag; Subsequently, the PA0644-PA0645 and PA0647-PA0648 gene subclones were inserted into the second cloning site of the pACYCDuet vector (between NdeI and XhoI restriction sites) without any additional tags.

### Protein purification and expression

Two types of tail-fibers were expressed in *E.coli* BL21 (DE3) cells. Strains harboring pETDuet-1 plasmids were cultured in fresh LB broth supplemented with 30 mg/L Ampicillin at 37 °C with shaking until reaching mid-log phase. Protein expression was induced with 0.2 mM isopropyl β-D-1-thiogalactopyranoside (IPTG), followed by overnight incubation at 16 °C or 28 °C. Cells were harvested by centrifugation, resuspended in lysis buffer (20 mM Tris-HCl, 150 mM NaCl, pH 8.0), and lysed via sonication on ice. The lysate was centrifuged at 10,000 × g for 40 min, and the clarified supernatant was loaded onto a Ni-NTA affinity column. After washing with a buffer containing 20 mM imidazole, bound proteins were eluted using stepwise gradients of 50 mM, 100 mM, and 500 mM imidazole. Target protein fractions were identified by SDS-PAGE, concentrated to 500 μL using a 3 kDa Amicon Ultra centrifugal filter, and further purified via size-exclusion chromatography on a Superdex 200 10/300 GL column pre-equilibrated with storage buffer (20 mM Tris-HCl, 150 mM NaCl, pH 8.0). Elution profiles were monitored at 220 nm and 280 nm, and fractions containing purified protein were pooled, concentrated again, and stored at −80°C.

### Proteomics analysis

The samples of F-pyocin from different mutation strains purified by size-exclusion chromatography were resuspended in lysis buffer containing 6 M guanidine hydrochloride, 10 mM tris (2-carboxyethyl) phosphine (TCEP), 40 mM chloroacetamide, and 100 mM Tris-HCl (pH 8.5), then heated at 95 °C for 10 minutes followed by 20 minutes of sonication to lyse the cells, reduce proteins, and perform alkylation. The resulting lysate was diluted 10-fold with the dilution buffer (25 mM Tris-HCl and 10% acetonitrile), and proteins were digested overnight at 37 °C with trypsin. Total peptides were acidified with 5% trifluoroacetic acid (TFA) and loaded onto C18 columns pre-equilibrated with methanol and 0.2% acetic acid. Columns were washed three times with 0.2% acetic acid and eluted with a buffer containing 80% acetonitrile and 0.2% acetic acid.

The samples were reconstituted in buffer A (0.1% formic acid, FA) and analyzed on an Orbitrap Fusion Lumos 480 mass spectrometer (Thermo Scientific) coupled online to an EASY-nLC 1200 system. A total of 0.5 µg of peptide digest was loaded onto a 28 cm-long, 75 µm inner diameter fused-silica capillary analytical column packed with 2 µm C18 reversed-phase particles. The mobile phases for liquid chromatography (LC) consisted of buffer A (0.1% FA) and buffer B (80% acetonitrile, 0.1% FA). Peptides were separated using a 65 min nonlinear gradient: 6-8% B (0-1 min), 8-25% B (1-44 min), 25-35% B (44-60 min), 35-55% B (60-61.5 min), 55-99% B (61.5-62 min), and 99% B (62-65 min), at a flow rate of 300 nL/min.

Mass spectrometric data were acquired in data-dependent acquisition (DDA) mode, with all measurements performed in the positive ion mode^49^. MS1 full scans were acquired at a resolution of 60,000 (at m/z 200), over a scan range of 350-1550 m/z, with an AGC target of 3.0 × 10⁶ and a maximum injection time of 50 ms. MS2 scans were acquired at a resolution of 15,000, using higher-energy collisional dissociation (HCD) with a normalized collision energy of 30%, an isolation window of 1.6 m/z, and a dynamic exclusion duration of 45 s. Data analysis was performed using pFind software^50^.

## Data Availability

The data that support this study are available from the corresponding authors upon request. The cryo-EM maps have been deposited in the Electron Microscopy Data Bank (EMDB) under accession codes EMD69233 (Cap); EMD69224 (Tube); EMD69223 (Tail-tip); and EMD69222 (Tail-fiber). The atomic coordinates have been deposited in the Protein Data Bank (PDB) under accession codes 23TF (Cap); 23SV (Tube); 23SU (Tail-tip); and 23ST (Tail-fiber).The mass spectrometry proteomics data have been deposited in the PRIDE partner repository of the ProteomeXchange Consortium under accession code PXD074579.

