## Supplemental Materials for "Structural basis for repurposing a flexible phage tail into an Intraspecific bacterial competition weapon"

**This PDF file includes:**

Supporting text

Figures S1 to S4

Tables S1 to S4

**Supporting Information Text**

**Figures**

**Figure S1** Single particle 3D reconstruction in the cap and tube regions of F-pyocin.

**A.** Representative cryo-EM micrograph of F-pyocin embedded in vitreous ice. The scale bar represents 50 nm. **B.** Flow chart for 3D reconstruction of the cap region in F-pyocin. **C.** Flow chart for 3D reconstruction of the tube region in F-pyocin. **D.** The fourier shell correlation (FSC) using the gold-standard criteria. **E.** Particle distribution orientation diagram. **F.** Left:cap region local resolution map (in Å); Right:tube region local resolution map (in Å).

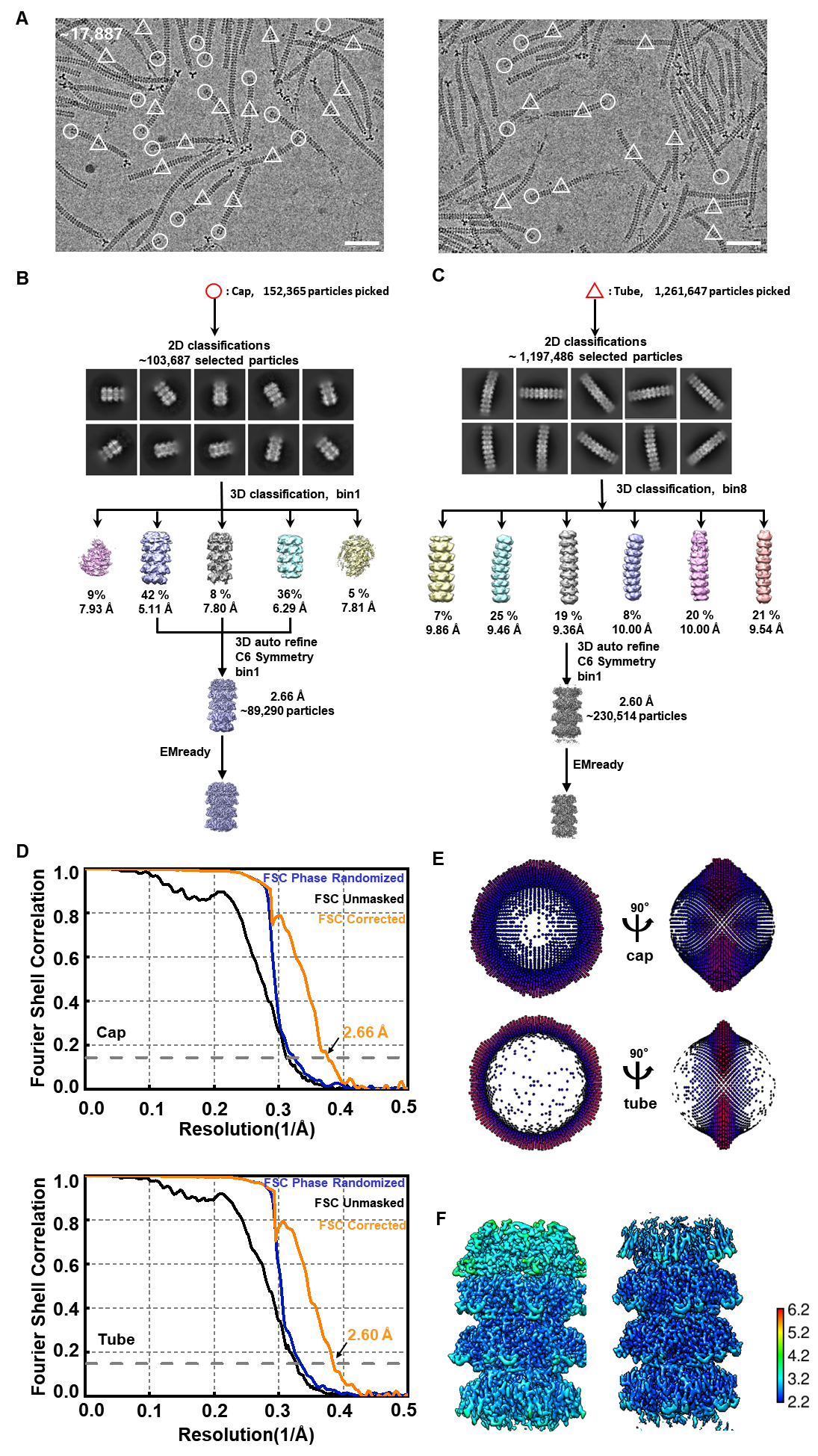

**Figure S2** Single particle 3D reconstruction in the tail tip and tail fiber regions of F-pyocin.

Figures **A, B, C, D,** and **E** are similar to supplementary Figure 1. **A.** The scale bar represents 50 nm. **E.** Left:tail tip region local resolution map (in Å); Right:tail fiber region local resolution map (in Å).

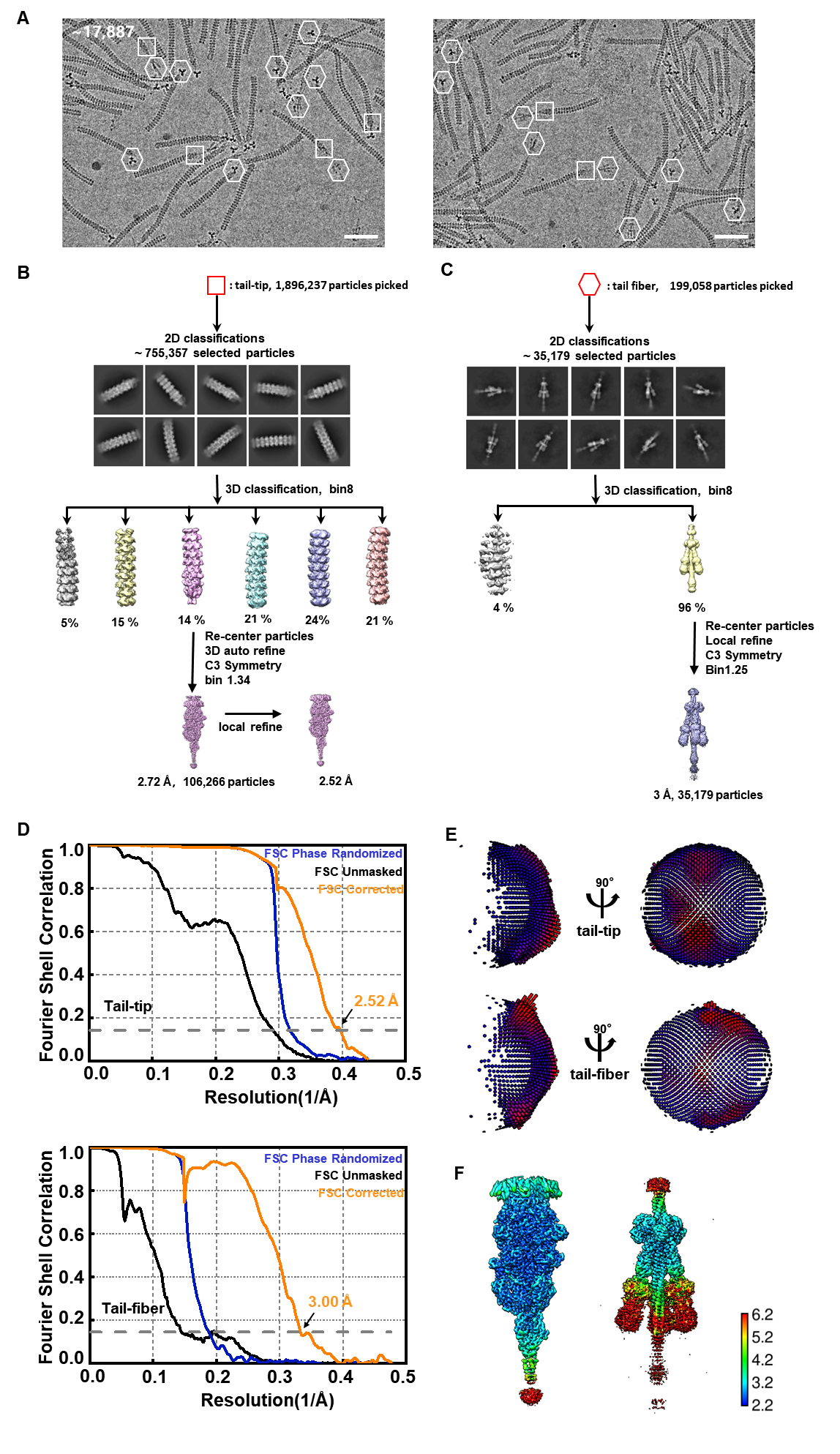

**Figure S3** Purification of recombinantly expressed full-length PA0640.

SDS-PAGE analysis of His-tagged PA0640 purified from *E. coli*. **M**, molecular weight marker; **L**, loading sample; **FT**, flow-through; **W**, wash (20 mM imidazole); **E1–E5**, elution fractions with increasing imidazole concentrations (50–500 mM). The major bands (dashed box) at ~15 kDa represent the C-terminal fragments of PA0640. The results indicate that full-length PA0640 is spontaneously processed by unknown host proteases, confirming the intrinsic instability of its N-terminal region.

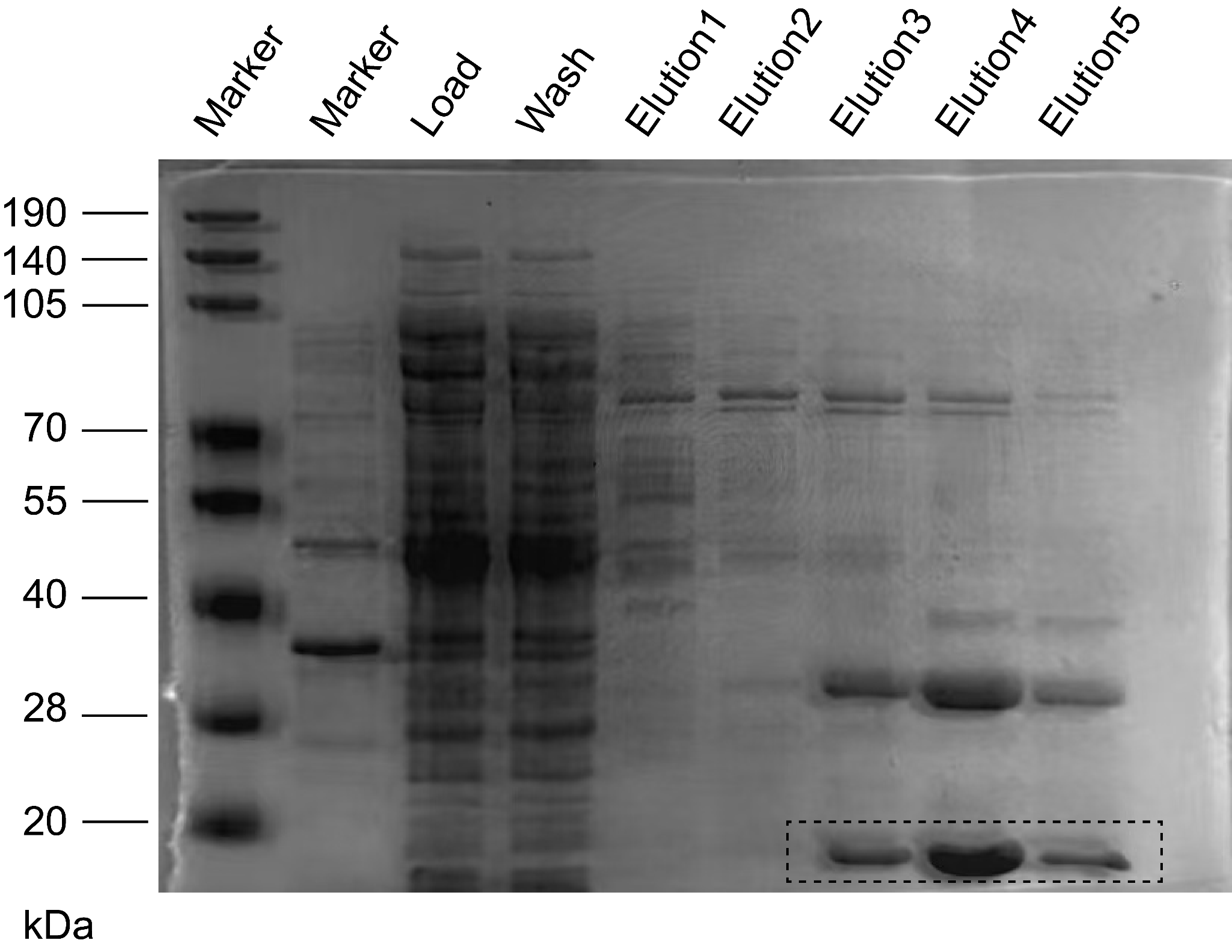

**Figure S4** Proteomic profiling of single-component deletion mutants of the F-pyocin.

Heatmap of F-pyocin components (y-axis) across WT and 17 knockout strains (x-axis) analyzed by DDA-MS. Color scale (blue to red) represents relative protein abundance. White boxes (NA) indicate non-detected proteins, highlighting assembly dependencies.

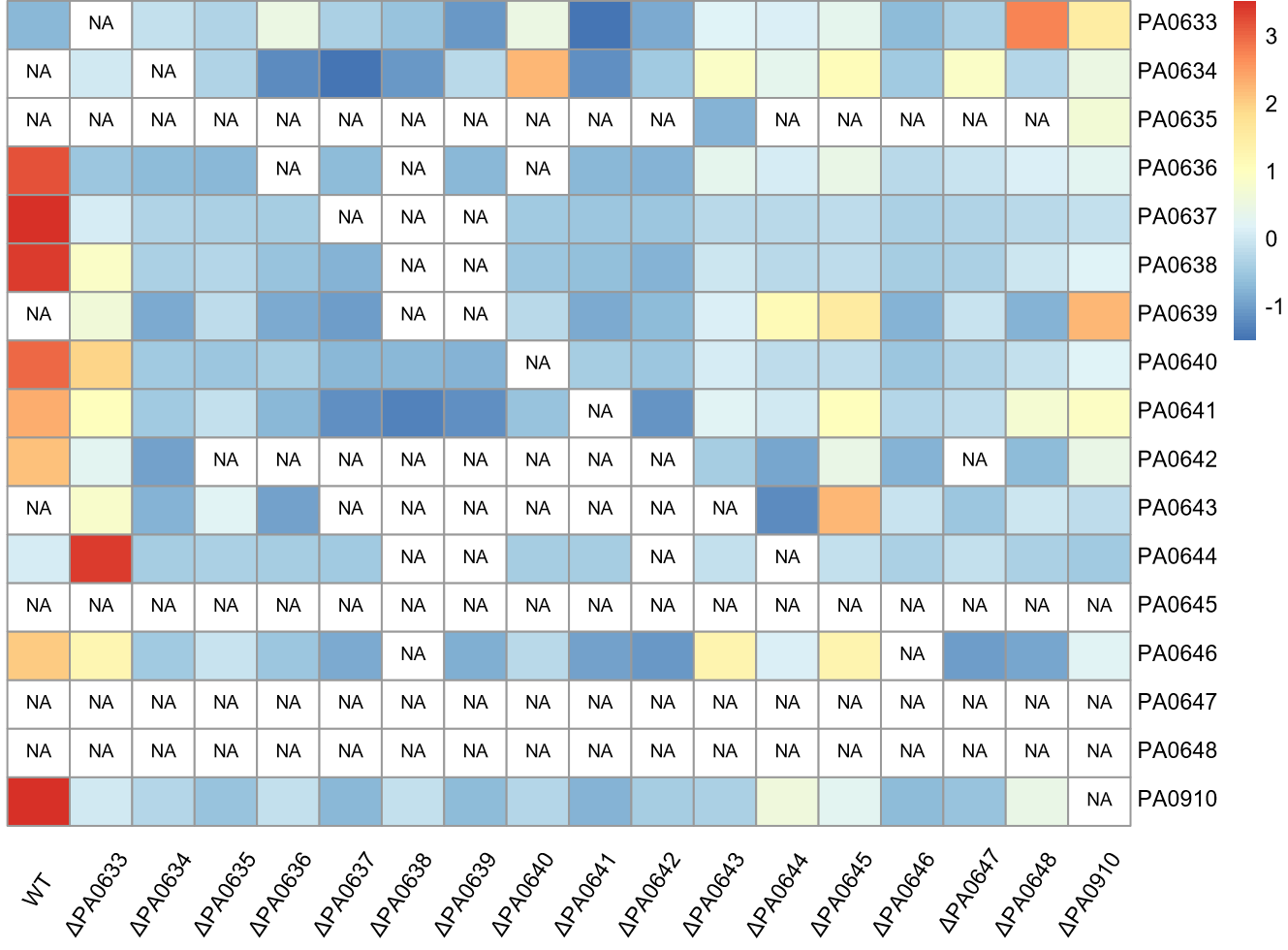

**Tables**

**Table S1** Cryo-EM data collection, refinement and validation statistics.

|  | **Cap**  **(EMDB ID 69233)**  **(PDB ID 23TF)** | **Tube**  **(EMDB ID 69224)**  **(PDB ID 23SV)** | **Tail-tip**  **(EMDB ID 69223)**  **(PDB ID 23SU)** | **Tail-fiber**  **(EMDB ID 69222)**  **(PDB ID 23ST)** |
| --- | --- | --- | --- | --- |
| **Data collection and processing** | | |  |  |
| Magnification | ×105,000 | ×105,000 | ×105,000 | ×105,000 |
| Voltage (kV) | 300 | 300 | 300 | 300 |
| Electron exposure (e-/Å2) | 60 | 60 | 60 | 60 |
| Defocus range tilted (μm) | -0.8~-2.2 | -0.8~-2.2 | -0.8~-2.2 | -0.8~-2.2 |
| Pixel size (Å) | 0.83 | 0.83 | 0.83 | 0.83 |
| Symmetry imposed | *C6* | *C6* | *C3* | *C3* |
| Initial particle images (no.) | 152,365 | 1,261,647 | 1,896,237 | 199,058 |
| Final particle images (no.) | 89,290 | 230,514 | 106,266 | 35,179 |
| Map resolution (Å) | 2.66 | 2.60 | 2.52 | 3.00 |
| FSC threshold | 0.143 | 0.143 | 0.143 | 0.143 |
| Map resolution range (Å) | 2.2-4.0 | 2.2-3.5 | 2.2-5.0 | 2.5-7.0 |
| Map sharpening B factor (Å2) | -96.2 | -96.4 | -78.5 | -62.2 |
| **Refinement** | | |  |  |
| Non hydrogen atoms | 14,178 | 14,760 | 41,310 | 29,919 |
| Protein/nucleotide residues | 1,854 | 1,956 | 5,361 | 4,035 |
| Ligands | / | / | SF4(3) | / |
| R.M.S.D. |  |  |  |  |
| Bond lengths (Å) | 0.11 | 0.11 | 0.10 | 0.13 |
| Bond angles (º) | 0.30 | 0.28 | 0.27 | 0.30 |
| **Validation** | | |  |  |
| MolProbity score | 1.48 | 1.29 | 1.73 | 2.01 |
| Clashscore | 6 | 6 | 8 | 12 |
| Rotamer outliers (%) | 0 | 0 | 0 | 0 |
| Ramachandran plot |  |  |  |  |
| Favored (%) | 97 | 98 | 96 | 94 |
| Allowed (%) | 3 | 2 | 4 | 6 |
| Outliers (%) | 0 | 0 | 0 | 0 |

**Table S2** Plasmids constructed in the experiments.

| Plasmid Number | Description | Resistance | Source |
| --- | --- | --- | --- |
| pK18mobSacB_ΔPA0622-0623 | Derivative of pK18mobsacB used for the in-frame deletion of *PA0622* and *PA0623* | GM | This paper |
| pK18mobSacB_ΔPA0633 | Derivative of pK18mobsacB used for the in-frame deletion of *PA0633* | GM | This paper |
| pK18mobSacB_ΔPA0634 | Derivative of pK18mobsacB used for the in-frame deletion of *PA0634* | GM | This paper |
| pK18mobSacB_ΔPA0635 | Derivative of pK18mobsacB used for the in-frame deletion of *PA0635* | GM | This paper |
| pK18mobSacB_ΔPA0636 | Derivative of pK18mobsacB used for the in-frame deletion of *PA0636* | GM | This paper |
| pK18mobSacB_ΔPA0637 | Derivative of pK18mobsacB used for the in-frame deletion of *PA0637* | GM | This paper |
| pK18mobSacB_ΔP0638 | Derivative of pK18mobsacB used for the in-frame deletion of *PA0638* | GM | This paper |
| pK18mobSacB_ΔP0639 | Derivative of pK18mobsacB used for the in-frame deletion of *PA0639* | GM | This paper |
| pK18mobSacB_ΔP0640 | Derivative of pK18mobsacB used for the in-frame deletion of *PA0640* | GM | This paper |
| pK18mobSacB_ΔP0641 | Derivative of pK18mobsacB used for the in-frame deletion of *PA0641* | GM | This paper |
| pK18mobSacB_ΔP0642 | Derivative of pK18mobsacB used for the in-frame deletion of *PA0642* | GM | This paper |
| pK18mobSacB_ΔP0643 | Derivative of pK18mobsacB used for the in-frame deletion of *PA0643* | GM | This paper |
| pK18mobSacB_ΔP0644 | Derivative of pK18mobsacB used for the in-frame deletion of *PA0644* | GM | This paper |
| pK18mobSacB_ΔP0645 | Derivative of pK18mobsacB used for the in-frame deletion of *PA0645* | GM | This paper |
| pK18mobSacB_ΔP0646 | Derivative of pK18mobsacB used for the in-frame deletion of *PA0646* | GM | This paper |
| pK18mobSacB_ΔP0647 | Derivative of pK18mobsacB used for the in-frame deletion of *PA0647* | GM | This paper |
| pK18mobSacB_ΔP0648 | Derivative of pK18mobsacB used for the in-frame deletion of *PA0648* | GM | This paper |
| pK18mobSacB_ΔP0910 | Derivative of pK18mobsacB used for the in-frame deletion of *PA0910* | GM | This paper |
| pETDuet_MCSǀ_PA0639 | PA0639 from *P. aeruginosa* PAO1 was cloned into MCS1 of pETDuet-1 under T7 promoter control. | Amp | This paper |
| pET28_PA0640 | PA0640 from *P. aeruginosa* PAO1 was cloned into pET-28a(+) under T7 promoter control. | Kan | This paper |
| pETDuet_MCS1_PA0643_MCS2_PA0644-PA0645 | *PA0643* from *P. aeruginosa* PAO1 was cloned into MCS1 of pETDuet-1 under T7 promoter control, while *PA0644* and *PA0645* were cloned into MCS2. Both were under control of T7 promoters. | Amp | This paper |
| pETDuet_MCS1_PA0646_MCS2_PA0647-PA0648 | *PA0646* from *P. aeruginosa* PAO1 was cloned into MCS1 of pETDuet-1 under T7 promoter control, while *PA0647* and *PA0648* were cloned into MCS2. Both were under control of T7 promoters. | Amp | This paper |

**Table S3** Bacteria strains in the experiments.

| Strains name | Description | Source |
| --- | --- | --- |
| E.coli DH5α | Used for constructing and amplifying plasmids | ATCC 68233 |
| E.coli BL21(DE3) | Used for heterologous protein expression under T7 promoter control | Laboratory  stock |
| E.coli WM3064 | Used as a donor strain for conjugal transfer of plasmids into *P. aeruginosa* PAO1 during gene knockout | Laboratory stock |
| Pseudomonas aeruginosa PAO1 | Used as the parental wild-type strain for the construction of mutant strains | ATCC 15692 |
| Pseudomonas aeruginosa JCM 5962 | This strain was used as the indicator strain for spot assays in this study | Lanzhou University Second Hospital |
| P.aeruginosa PAO1 ΔPA0622-PA0623 | Mutant strain with PA0622 and PA0623 knocked out in the coding frame | This paper |
| P.aeruginosa PAO1 ΔPA0633 | Mutant strain with PA0633 knocked out in the coding frame | This paper |
| P.aeruginosa PAO1 ΔPA0634 | Mutant strain with PA0634 knocked out in the coding frame | This paper |
| P.aeruginosa PAO1 ΔPA0635 | Mutant strain with PA0635 knocked out in the coding frame | This paper |
| P.aeruginosa PAO1 ΔPA0636 | Mutant strain with PA0636 knocked out in the coding frame | This paper |
| P.aeruginosa PAO1 ΔPA0637 | Mutant strain with PA0637 knocked out in the coding frame | This paper |
| P.aeruginosa PAO1 ΔPA0638 | Mutant strain with PA0638 knocked out in the coding frame | This paper |
| P.aeruginosa PAO1 ΔPA0639 | Mutant strain with PA0639 knocked out in the coding frame | This paper |
| P.aeruginosa PAO1 ΔPA0640 | Mutant strain with PA0640 knocked out in the coding frame | This paper |
| P.aeruginosa PAO1 ΔPA0641 | Mutant strain with PA0641 knocked out in the coding frame | This paper |
| P.aeruginosa PAO1 ΔPA0642 | Mutant strain with PA0642 knocked out in the coding frame | This paper |
| P.aeruginosa PAO1 ΔPA0643 | Mutant strain with PA0643 knocked out in the coding frame | This paper |
| P.aeruginosa PAO1 ΔPA0644 | Mutant strain with PA0644 knocked out in the coding frame | This paper |
| P.aeruginosa PAO1 ΔPA0645 | Mutant strain with PA0633 knocked out in the coding frame | This paper |
| P.aeruginosa PAO1 ΔPA0646 | Mutant strain with PA0633 knocked out in the coding frame | This paper |
| P.aeruginosa PAO1 ΔPA0647 | Mutant strain with PA0633 knocked out in the coding frame | This paper |
| P.aeruginosa PAO1 ΔPA0648 | Mutant strain with PA0633 knocked out in the coding frame | This paper |
| P.aeruginosa PAO1 ΔPA0910 | Mutant strain with PA0633 knocked out in the coding frame | This paper |

**Table S4** Primers used in this study.

| **Name** | **Description** | **Sequence** |
| --- | --- | --- |
| pK18mobSacB_ΔPA0622-0623_Up_F | For construction of the plasmids  pK18mobSacB_ΔPA0622-0623 | GCTATGACATGATTACGAATTCCGCACATAACCATGACGTCGGACAGCT |
| pK18mobSacB_ΔPA0622-0623_Up_R | For construction of the plasmids  pK18mobSacB_ΔPA0622-0623 | CGTGAACACATCGCAGAGGCCGATGAC |
| pK18mobSacB_ΔPA0622-0623_Down_F | For construction of the plasmids  pK18mobSacB_ΔPA0622-0623 | GCCTCTGCGATGTGTTCACGGTTGCGGTCAGCTACTACAAGCTGGAAGTCGAT |
| pK18mobSacB_ΔPA0622-0623_Down_R | For construction of the plasmids  pK18mobSacB_ΔPA0622-0623 | TTTCCACGGTGTGCGTCTAGAGGCATTGCCTGGCTGGCCTGCTC |
| pK18mobSacB_ΔPA0622-0623_F | For construction of the plasmids  pK18mobSacB_ΔPA0622-0623 | GACGCCCGTCCGTACCAGATTGGCTC |
| pK18mobSacB_ΔPA0622-0623_R | For construction of the plasmids  pK18mobSacB_ΔPA0622-0623 | GCACTGTGCCGGACGCGAGAGAC |
| pK18mobSacB_ΔPA0633_Up_F | For construction of the plasmids  pK18mobSacB_ΔPA0633 | GCTATGACATGATTACGAATTCATGAGCCGGCTCGCTCTGCTCCT |
| pK18mobSacB_ΔPA0633_Up_R | For construction of the plasmids  pK18mobSacB_ΔPA0633 | CGGGTTGAACGAGGTCACGCCTTCGAT |
| pK18mobSacB_ΔPA0633_Down_F | For construction of the plasmids  pK18mobSacB_ΔPA0633 | GCGTGACCTCGTTCAACCCGGAAGGCTACGTCAGCGACTTCCCCTTCGAT |
| pK18mobSacB_ΔPA0633_Down_R | For construction of the plasmids  pK18mobSacB_ΔPA0633 | TTTCCACGGTGTGCGTCTAGATGTCCAGCGCCTGGTCCAGCACG |
| pK18mobSacB_ΔPA0633_F | For construction of the plasmids  pK18mobSacB_ΔPA0633 | CACGGCTGACCGTCCTGGAGGC |
| pK18mobSacB_ΔPA0633_R | For construction of the plasmids  pK18mobSacB_ΔPA0633 | GGAACGCGGGTCCAGGTAATCTCCTTG |
| pK18mobSacB_ΔPA0634_Up_F | For construction of the plasmids  pK18mobSacB_ΔPA0634 | GCTATGACATGATTACGAATTCCTGAAGGCCGGCGATTCGCTGC |
| pK18mobSacB_ΔPA0634_Up_R | For construction of the plasmids  pK18mobSacB_ΔPA0634 | CCAGGTAATCTCCTTGCGCACCAGC |
| pK18mobSacB_ΔPA0634_Down_F | For construction of the plasmids  pK18mobSacB_ΔPA0634 | CGCAAGGAGATTACCTGGGCGCTGGGCTTCCTGCTGCTG |
| pK18mobSacB_ΔPA0634_Down_R | For construction of the plasmids  pK18mobSacB_ΔPA0634 | TTTCCACGGTGTGCGTCTAGAGTCGCCAGCCTTCGCCTGTCC |
| pK18mobSacB_ΔPA0634_F | For construction of the plasmids  pK18mobSacB_ΔPA0634 | TCCGGCGCCGGCAAGTGGAC |
| pK18mobSacB_ΔPA0634_R | For construction of the plasmids  pK18mobSacB_ΔPA0634 | CGACCCGCCGACGCCGTTCAG |
| pK18mobSacB_ΔPA0635_Up_F | For construction of the plasmids  pK18mobSacB_ΔPA0635 | GCTATGACATGATTACGAATTCTCCTGGAGGCGGGGCCAGAC |
| pK18mobSacB_ΔPA0635_Up_R | For construction of the plasmids  pK18mobSacB_ΔPA0635 | CCGCTCCTTGGCCTCGGCAAT |
| pK18mobSacB_ΔPA0635_Down_F | For construction of the plasmids  pK18mobSacB_ΔPA0635 | TTGCCGAGGCCAAGGAGCGGATGGCTCTCCAGCCCCTTTCGCT |
| pK18mobSacB_ΔPA0635_Down_R | For construction of the plasmids  pK18mobSacB_ΔPA0635 | TTTCCACGGTGTGCGTCTAGAGATGCCGGCCTTGCCATTCATCCAGC |
| pK18mobSacB_ΔPA0635_F | For construction of the plasmids  pK18mobSacB_ΔPA0635 | GCTGCAGGCGGTCGGTGAGG |
| pK18mobSacB_ΔPA0635_R | For construction of the plasmids  pK18mobSacB_ΔPA0635 | GCGTCAGGCTGCCGTCGGTAT |
| pK18mobSacB_ΔPA0636_Up_F | For construction of the plasmids  pK18mobSacB_ΔPA0636 | GCTATGACATGATTACGAATTCCGAGCGTCAAGTGGGCCATCGG |
| pK18mobSacB_ΔPA0636_Up_R | For construction of the plasmids  pK18mobSacB_ΔPA0636 | CTCCATGGCGCGCATTTTCTGATCGTT |
| pK18mobSacB_ΔPA0636_Down_F | For construction of the plasmids  pK18mobSacB_ΔPA0636 | CAGAAAATGCGCGCCATGGAGAGCGGCGACGACGGTACGGGAC |
| pK18mobSacB_ΔPA0636_Down_R | For construction of the plasmids  pK18mobSacB_ΔPA0636 | TTTCCACGGTGTGCGTCTAGATCGAACATCGCCGTACCGGTGTAGTTGCAG |
| pK18mobSacB_ΔPA0636_F | For construction of the plasmids  pK18mobSacB_ΔPA0636 | TGGCTCTCCAGCCCCTTTCGCTGC |
| pK18mobSacB_ΔPA0636_R | For construction of the plasmids  pK18mobSacB_ΔPA0636 | GGCTGTAGCCGCCGCCGTACT |
| pK18mobSacB_ΔPA0637_Up_F | For construction of the plasmids  pK18mobSacB_ΔPA0637 | GCTATGACATGATTACGAATTC  CGGCATCACCGAGGCGCAACT |
| pK18mobSacB_ΔPA0637_Up_R | For construction of the plasmids  pK18mobSacB_ΔPA0637 | CTGCACCTGACGTACCAGTTGGTTCGCCTG |
| pK18mobSacB_ΔPA0637_Down_F | For construction of the plasmids  pK18mobSacB_ΔPA0637 | CTGGTACGTCAGGTGCAGCAACGGCGTGTTCACCCTGAACACCACC |
| pK18mobSacB_ΔPA0637_Down_R | For construction of the plasmids  pK18mobSacB_ΔPA0637 | TTTCCACGGTGTGCGTCTAGACAGACCATGCAGTTCGCAACTGACCCGAT |
| pK18mobSacB_ΔPA0637_F | For construction of the plasmids  pK18mobSacB_ΔPA0637 | CGCAACGTGGTGGCGCAGGAG |
| pK18mobSacB_ΔPA0637_R | For construction of the plasmids  pK18mobSacB_ΔPA0637 | CAGCGCACCAGGCTCGAGGGT |
| pK18mobSacB_ΔPA0638_Up_F | For construction of the plasmids  pK18mobSacB_ΔPA0638 | GCTATGACATGATTACGAATTCTC  GCCGACTCGGTGATCCAGGATAT |
| pK18mobSacB_ΔPA0638_Up_R | For construction of the plasmids  pK18mobSacB_ΔPA0638 | CCCCTGCTGCAGGTGGCCGTG |
| pK18mobSacB_ΔPA0638_Down_F | For construction of the plasmids  pK18mobSacB_ΔPA0638 | CACGGCCACCTGCAGCAGGGGGT  GGACGATCCGGCGCTGGACC |
| pK18mobSacB_ΔPA0638_Down_R | For construction of the plasmids  pK18mobSacB_ΔPA0638 | TTTCCACGGTGTGCGTCTAGAGAAGCGCTTGCCCAGAACCCCGTAC |
| pK18mobSacB_ΔPA0638_F | For construction of the plasmids  pK18mobSacB_ΔPA0638 | CACCCTGAACACCACCTTCCAGCAAGT |
| pK18mobSacB_ΔPA0638_R | For construction of the plasmids  pK18mobSacB_ΔPA0638 | CGATGGCCCGCTGCAGGCT |
| pK18mobSacB_ΔPA0639_Up_F | For construction of the plasmids  pK18mobSacB_ΔPA0639 | GCTATGACATGATTACGAATTCGCTGATCGGCCCGATCCGCGATT |
| pK18mobSacB_ΔPA0639_Up_R | For construction of the plasmids  pK18mobSacB_ΔPA0639 | GGCCACGTAGCGGCGTTGACGCAC |
| pK18mobSacB_ΔPA0639_Down_F | For construction of the plasmids  pK18mobSacB_ΔPA0639 | CAACGCCGCTACGTGGCCGACGGACCGTTCCTGCTGCACCAC |
| pK18mobSacB_ΔPA0639_Down_R | For construction of the plasmids  pK18mobSacB_ΔPA0639 | TTTCCACGGTGTGCGTCTAGAGCCAGCAGCGGCGTATCGTCCAG |
| pK18mobSacB_ΔPA0639_F | For construction of the plasmids  pK18mobSacB_ΔPA0639 | CGCTGTCCCACGGCGGCTTC |
| pK18mobSacB_ΔPA0639_R | For construction of the plasmids  pK18mobSacB_ΔPA0639 | CTGTTCCTCTCGTGAAGAGCCGATTCCGAC |
| pK18mobSacB_ΔPA0640_Up_F | For construction of the plasmids  pK18mobSacB_ΔPA0640 | GCTATGACATGATTACGAATTCCTGCGCTTCGGCGCGGACAAC |
| pK18mobSacB_ΔPA0640_Up_R | For construction of the plasmids  pK18mobSacB_ΔPA0640 | CTCGCGTACCGTGCCGCTTTCCAACAATCG |
| pK18mobSacB_ΔPA0640_Down_F | For construction of the plasmids  pK18mobSacB_ΔPA0640 | GAAAGCGGCACGGTACGCGAGGCCTTCGGCGGCCCGGTC |
| pK18mobSacB_ΔPA06490_Down_R | For construction of the plasmids  pK18mobSacB_ΔPA0640 | TTTCCACGGTGTGCGTCTAGAGGGTTGCTGGTCCAGGCCGACTTG |
| pK18mobSacB_ΔPA0640_F | For construction of the plasmids  pK18mobSacB_ΔPA0640 | GCGAGCCGTCGGAATCGGCTCTT |
| pK18mobSacB_ΔPA0640_R | For construction of the plasmids  pK18mobSacB_ΔPA0640 | CATGGCGGGCCTGTTACGGCGTT |
| pK18mobSacB_ΔPA0641_Up_F | For construction of the plasmids  pK18mobSacB_ΔPA0641 | GCTATGACATGATTACGAATTCGACCGCTCTTGAGGGCAAGACCACGC |
| pK18mobSacB_ΔPA0641_Up_R | For construction of the plasmids  pK18mobSacB_ΔPA0641 | TCCGTCCTGAATCAGGGCGTTCTTGATAAACATCTGTCC |
| pK18mobSacB_ΔPA0641_Down_F | For construction of the plasmids  pK18mobSacB_ΔPA0641 | AACGCCCTGATTCAGGACGGATGAGTATCTATGGATTACGTATCTATCGAGAGAATGGGACCGCCG |
| pK18mobSacB_ΔPA0641_Down_R | For construction of the plasmids  pK18mobSacB_ΔPA0641 | TTTCCACGGTGTGCGTCTAGATCGAGGACACCGCCCAGGGTTTGATCC |
| pK18mobSacB_ΔPA0641_F | For construction of the plasmids  pK18mobSacB_ΔPA0641 | TCGGTTTATCTGGTTCGACAGCTCCAGCGG |
| pK18mobSacB_ΔPA0641_R | For construction of the plasmids  pK18mobSacB_ΔPA0641 | GCATCAACCCGTCTGGAATATACAACCCGCAACG |
| pK18mobSacB_ΔPA0642_Up_F | For construction of the plasmids  pK18mobSacB_ΔPA0642 | GCTATGACATGATTACGAATTCGCAGAAGCAGGTCGGCGACCTCG |
| pK18mobSacB_ΔPA0642_Up_R | For construction of the plasmids  pK18mobSacB_ΔPA0642 | TACAGTCTGCCCCAAGCCATTCCTTACCAAGACC |
| pK18mobSacB_ΔPA0642_Down_F | For construction of the plasmids  pK18mobSacB_ΔPA0642 | AATGGCTTGGGGCAGACTGTAAGGCTTATGCAGCATGCATATACCGGGCT |
| pK18mobSacB_ΔPA0642_Down_R | For construction of the plasmids  pK18mobSacB_ΔPA0642 | TTTCCACGGTGTGCGTCTAGAATGCCTCCGGAGGCAGGTACCCC |
| pK18mobSacB_ΔPA0642_F | For construction of the plasmids  pK18mobSacB_ΔPA0642 | TGGGCCCGAATGGGTGATGGCGTT |
| pK18mobSacB_ΔPA0642_R | For construction of the plasmids  pK18mobSacB_ΔPA0642 | AACCGAACCTTTCGAATGCCAAGCCATAGCAT |
| pK18mobSacB_ΔPA0643_Up_F | For construction of the plasmids  pK18mobSacB_ΔPA0643 | GCTATGACATGATTACGAATTCCGACAGCTCCAGCGGGACGGC |
| pK18mobSacB_ΔPA0643_Up_R | For construction of the plasmids  pK18mobSacB_ΔPA0643 | GCCACGGAACGCATCGCCGGTTC |
| pK18mobSacB_ΔPA0643_Down_F | For construction of the plasmids  pK18mobSacB_ΔPA0643 | ACCGGCGATGCGTTCCGTGGCCGTCCCTGGGGTTCGACGATGTCGG |
| pK18mobSacB_ΔPA0643_Down_R | For construction of the plasmids  pK18mobSacB_ΔPA0643 | TTTCCACGGTGTGCGTCTAGAGCGGGTACTGGCGTCGAATACCATCGACAG |
| pK18mobSacB_ΔPA0644_Up_F | For construction of the plasmids  pK18mobSacB_ΔPA0644 | GCTATGACATGATTACGAATTCGCCAAGGCTTATGCAGCATGCATATACCGGG |
| pK18mobSacB_ΔPA0644_Up_R | For construction of the plasmids  pK18mobSacB_ΔPA0644 | GCTGAGACGTTCCCTGTCAGGCGTGATGC |
| pK18mobSacB_ΔPA0644_Down_F | For construction of the plasmids  pK18mobSacB_ΔPA0644 | CCTGACAGGGAACGTCTCAGCCCGGTCGACCAGGATGGAAAAGTCGAGGCGTCAGCGACTTCCCCTTCGAT |
| pK18mobSacB_ΔPA0644_Down_R | For construction of the plasmids  pK18mobSacB_ΔPA0644 | TTTCCACGGTGTGCGTCTAGAGAAGCCCGTAACGAAGGCAGCTGGTAGC |
| pK18mobSacB_ΔPA0644_F | For construction of the plasmids  pK18mobSacB_ΔPA0644 | CACCGCCAGCGTGGCCAGC |
| pK18mobSacB_ΔPA0644_R | For construction of the plasmids  pK18mobSacB_ΔPA0644 | CCAAGGCCTGATCGCGCACATCTGC |
| pK18mobSacB_ΔPA0645_Up_F | For construction of the plasmids  pK18mobSacB_ΔPA0645 | GCTATGACATGATTACGAATTCTGGCCCTGACGGCCGGTGGTAC |
| pK18mobSacB_ΔPA0645_Up_R | For construction of the plasmids  pK18mobSacB_ΔPA0645 | GGCCTTTGCAGGGCTGACGTACTTATCCCAT |
| pK18mobSacB_ΔPA0645_Down_F | For construction of the plasmids  pK18mobSacB_ΔPA0645 | TACGTCAGCCCTGCAAAGGCCGCCAAGGTGGAAGAGATCAAGTCACGATACCCGA |
| pK18mobSacB_ΔPA0645_Down_R | For construction of the plasmids  pK18mobSacB_ΔPA0645 | TTTCCACGGTGTGCGTCTAGATGGCAGAGGCGGGCAGATATCCCCC |
| pK18mobSacB_ΔPA0645_F | For construction of the plasmids  pK18mobSacB_ΔPA0645 | CCAGGGAGATCGAGTCAATTCCGGTCGACCAG |
| pK18mobSacB_ΔPA0645_R | For construction of the plasmids  pK18mobSacB_ΔPA0645 | GGGTTAGTTCGGAACCACAGGGCTGACGC |
| pK18mobSacB_ΔPA0646_Up_F | For construction of the plasmids  pK18mobSacB_ΔPA0646 | GCTATGACATGATTACGAATTCTTCCGCCACTGGCCAGTACATGCG |
| pK18mobSacB_ΔPA0646_Up_R | For construction of the plasmids  pK18mobSacB_ΔPA0646 | CGGCCCTCTGAAGGCATCGCCG |
| pK18mobSacB_ΔPA0646_Down_F | For construction of the plasmids  pK18mobSacB_ΔPA0646 | GGCGATGCCTTCAGAGGGCCGGGGGTTCGTGGGGCCTATCTCAACGGT |
| pK18mobSacB_ΔPA0646_Down_R | For construction of the plasmids  pK18mobSacB_ΔPA0646 | TTTCCACGGTGTGCGTCTAGAGGGAAAGGACGATGCGCTCGGGC |
| pK18mobSacB_ΔPA0646_F | For construction of the plasmids  pK18mobSacB_ΔPA0646 | CCCTATCAAGCCCGCCGACTGCGG |
| pK18mobSacB_ΔPA0646_R | For construction of the plasmids  pK18mobSacB_ΔPA0646 | CGAGTCCTCGGGCAGGACTACAACCG |
| pK18mobSacB_ΔPA0647_Up_F | For construction of the plasmids  pK18mobSacB_ΔPA0647 | GCTATGACATGATTACGAATTCCCCTATCAAGCCCGCCGACTGCGG |
| pK18mobSacB_ΔPA0647_Up_R | For construction of the plasmids  pK18mobSacB_ΔPA0647 | GACATCCATATCAGTCGAGTCCTCGGGCAGGA |
| pK18mobSacB_ΔPA0647_Down_F | For construction of the plasmids  pK18mobSacB_ΔPA0647 | GACTCGACTGATATGGATGTCTTCCCTGTCCCCATCCATATCGTTGAAGACGGA |
| pK18mobSacB_ΔPA0647_Down_R | For construction of the plasmids  pK18mobSacB_ΔPA0647 | TTTCCACGGTGTGCGTCTAGATCGTCGAGCGATCCGTCGGCATGT |
| pK18mobSacB_ΔPA0647_F | For construction of the plasmids  pK18mobSacB_ΔPA0647 | GGCTAACGACACCATTACGGTGATGGCTGTGG |
| pK18mobSacB_ΔPA0647_R | For construction of the plasmids  pK18mobSacB_ΔPA0647 | CCGCTCTCTGGAAACCTGATCCGGTGT |
| pK18mobSacB_ΔPA0648_Up_F | For construction of the plasmids  pK18mobSacB_ΔPA0648 | GCTATGACATGATTACGAATTCGGGTTCCCTCCTGGCGGCAGT |
| pK18mobSacB_ΔPA0648_Up_R | For construction of the plasmids  pK18mobSacB_ΔPA0648 | CCCGCTCTCTGGAAACCTGATCCGGTG |
| pK18mobSacB_ΔPA0648_Down_F | For construction of the plasmids  pK18mobSacB_ΔPA0648 | ATCAGGTTTCCAGAGAGCGGGAGATATCCGCTCCCGCGCATCTGACCT |
| pK18mobSacB_ΔPA0648_Down_R | For construction of the plasmids  pK18mobSacB_ΔPA0648 | TTTCCACGGTGTGCGTCTAGATCCTCGGTGGTCAGGTCGAGCTGGTT |
| pK18mobSacB_ΔPA0648_F | For construction of the plasmids  pK18mobSacB_ΔPA0648 | GGAGCTACAGCGAAGCTGCCAAATTCCCT |
| pK18mobSacB_ΔPA0648_R | For construction of the plasmids  pK18mobSacB_ΔPA0648 | CCCGACAGTAGCGATCTAAAATGGCGAAGCCT |
| pK18mobSacB_ΔPA0910_Up_F | For construction of the plasmids  pK18mobSacB_ΔPA0910 | GCTATGACATGATTACGAATTCGCTGTCGAAGAGCACCCTGCATCGCTG |
| pK18mobSacB_ΔPA0910_Up_R | For construction of the plasmids  pK18mobSacB_ΔPA0910 | CAGCAGTTGATCGGCATCGATGACGCTGC |
| pK18mobSacB_ΔPA0910_Down_F | For construction of the plasmids  pK18mobSacB_ΔPA0910 | ATCGATGCCGATCAACTGCTGCTTGTTCTCGCTCACGAACACGTGCTGCT |
| pK18mobSacB_ΔPA0910_Down_R | For construction of the plasmids  pK18mobSacB_ΔPA0910 | TTTCCACGGTGTGCGTCTAGAGTCCCTCCTTCTCCTCCTCTCGCTGGCGTCCAGCGCCTGGTCCAGCACG |
| pK18mobSacB_ΔPA0910_F | For construction of the plasmids  pK18mobSacB_ΔPA0910 | GTCGGTGCCTGTCCGTCATCCATCCATCC |
| pK18mobSacB_ΔPA0910_R | For construction of the plasmids  pK18mobSacB_ΔPA0910 | GGATCTCTCCTCTATGTATGGATGGCATCGCTACGATGC |
| pETDuet_MCSǀ_PA0639_F | For construction of the plasmid pETDuet_MCSǀ_PA0639 | CATCATCACCACAGCCAGGATCCGATGGAACTGAGCCGCAGCCT |
| pETDuet_MCSǀ_PA0639_R | For construction of the plasmid pETDuet_MCSǀ_PA0639 | TAAGCATTATGCGGCCGCAAGCTTTCACTGCGGCATCCGCGTG |
| pET28_PA0640-F | For construction of the plasmid pET28_PA0640 | GTTTAACTTTAAGAAGGAGATATACCATGGGAATGAGTGACACCCTGAGTCAGGGC |
| pET28_PA0640-R | For construction of the plasmid pET28_PA0640 | TGGTGGTGGTGGTGGTGCTCGAGCAGCCGGTCCTCGGCATAGA |
| pETDuet_PA0643_F | For construction of the plasmid pETDuet_MCS1_PA0643_MCS2_PA0644-PA0645 | CATCATCACCACAGCCAGGATCCAATGGCTTGGCATTCGAAAGGTTCGG |
| pETDuet_PA0643_R | For construction of the plasmid pETDuet_MCS1_PA0643_MCS2_PA0644-PA0645 | TAAGCATTATGCGGCCGCAAGCTTTCAATTCCACCTCCCCATGGCGACA |
| pETDuet_PA0644-0645_F | For construction of the plasmid pETDuet_MCS1_PA0643_MCS2_PA0644-PA0645 | GTTAAGTATAAGAAGGAGATATACATATGATGAAGCTGTTGTTGAAACCTTTGTTGCAGCAG |
| pETDuet_PA0644-0645_R | For construction of the plasmid  pETDuet_MCS1_PA0646_MCS2_PA0647-PA0648 | GCGGTTTCTTTACCAGACTCGAGCTATTCAGGCATCGGGTATCGTGACTTGATCTCT |
| pETDuet_PA0646_F | For construction of the plasmid  pETDuet_MCS1_PA0646_MCS2_PA0647-PA0648 | CATCATCACCACAGCCAGGATCCAATGCCTTGGTATTCCACAGGCACGG |
| pETDuet_PA0646_R | For construction of the plasmid  pETDuet_MCS1_PA0646_MCS2_PA0647-PA0648 | TAAGCATTATGCGGCCGCAAGCTTTCAGTACCACCTCCCCACAGCCA |
| pETDuet_PA0647-PA0648_F | For construction of the plasmid  pETDuet_MCS1_PA0646_MCS2_PA0647-PA0648 | GTTAAGTATAAGAAGGAGATATACATATGATGCGTATCGAACTCAGTCCGGTTGTAGT |
| pETDuet_PA0647-PA0648_R | For construction of the plasmid  pETDuet_MCS1_PA0646_MCS2_PA0647-PA0648 | GCGGTTTCTTTACCAGACTCGAGTCAGATGCGCGGGAGCGGATATCTAGC |
